# Quantifying the diverse contributions of hierarchical muscle interactions to motor function

**DOI:** 10.1101/2023.11.30.569159

**Authors:** David O’Reilly, William Shaw, Pauline Hilt, Rafael de Castro Aguiar, Sarah L Astill, Ioannis Delis

## Abstract

The muscle synergy concept suggests that the human motor system is organised into functional modules comprised of muscles *‘working together’* towards common task-goals. This study offers a nuanced computational perspective to muscle synergies, where muscles interacting across multiple scales have functionally-similar, - complementary and -independent roles. Making this viewpoint implicit to a methodological approach applying Partial Information Decomposition to large-scale muscle activations, we unveiled nested networks of functionally diverse inter- and intra-muscular interactions with distinct functional consequences on task performance. This approach’s effectiveness is demonstrated using simulations and by extracting generalisable muscle networks from benchmark datasets of muscle activity. Specific network components are shown to correlate with a) balance performance and b) differences in motor variability between young and older adults. By aligning muscle synergy analysis with leading theoretical insights on movement modularity, the mechanistic insights presented here suggest the proposed methodology offers enhanced research opportunities towards health and engineering applications.

## Introduction

Hierarchical modularity is a ubiquitous characteristic of complex living systems such as the human nervous system ^1,2^. The constituent parts (i.e. individual neurons, muscles etc.) at each level interact in a goal-directed manner ^3^, forming functionally specialized modules that cooperate towards common task-goals. The putative interactions within these modules include a common selectivity (i.e. redundancy) for task information along with unique contributions by individual parts to the processing of task information. To exemplify this point in the context of motor control, in seminal work Ivanenko and colleagues noted: “*…We can think of (*electromyographic*) EMG waveforms as being dependent on two aspects. First, there are some underlying common waveforms shared by the muscles. Second, each muscle also captures a unique aspect of activation that is not addressed by any other muscle”* ^4^. However, for a set of muscle activity patterns to form an emergent whole (i.e. a coordinated movement), the integration of redundant and unique muscle constituents in the form of cross-module, synergistic muscle interactions is necessary. Synergistic interactions combine information across functionally heterogeneous modules, therefore serving as important channels of communication for the integration of information in the nervous system ^5,6^. The complementary information generated by these interactions is super-additive, emerging from the union of lower-level constituents ^7^. Indeed, from a more coarse-grained view, these emergent functional modules themselves, through hierarchies of complementary interactions ^8^, form parts of greater wholes (i.e. submodules-within-modules) ^9^.

Motor control research, has focused primarily on deciphering how the numerous degrees-of-freedom of the human body are coordinated for everyday tasks ^10,11^. This avenue of research uses the ‘*muscle synergy’* as a guiding concept ^12,13^, where the cohesive interactions between groups of muscles (‘*muscle synergies’*) map to common task-goals, and in doing so simplify movement execution. This definition of synergy hinges on covariations between muscles that represent their functional cooperation (‘*working together’*) and is distinct from the information-theoretic based description we have previously provided here. To avoid confusion in the use of the synergy term, here we refer to muscle covariations as couplings or interactions and, based on our framework presented below, we separate these further into “redundant” (i.e. functionally similar) and “synergistic” (i.e. functionally complementary) types of interaction. Thus, unlike the traditional definition of synergy in motor control ^12^, here synergy is a specific type of muscle interaction where muscles cooperate towards different, complementary aspects of task performance. This distinction in muscle interaction types is inspired by recent influential works suggesting that a more complex functional architecture underlies human motor control ^14–17^. For instance, anatomically proximal musculature thought to have equivalent task efficacy have, in fact, demonstrated a partial decoupling and sharing of common drive with distal musculature ^14,15^. This distributed neural architecture strongly suggests that muscle interactions contribute in functionally diverse ways to task performance. Moreover, the identification of independent functional modules indicates that multiple sources of common drive are likely present in the muscle space ^17^. Altogether, this recent perspective proposes that functional modularity exists both between and within muscles, simplifying the control of movement by enabling their flexible compliance towards multiple task-objectives ^18^. It also integrates independent muscle control as a fine-grained control mechanism into the conceptual perspectives on course-grained motor control, importantly broadening the context of human movement modularity for the research field. It is therefore prescient for the field to develop analytical approaches to comprehensively understand human movement modularity from this nuanced perspective. Hence, an objective of this study is to specifically address this research gap by providing a bridge between theoretical and computational frameworks within the motor control field.

In ^19^, we addressed this research gap by proposing a computational framework that dissects task information into task-irrelevant (i.e. present across all tasks), -redundant and -synergistic muscular interactions. In doing so, we aligned current analytical approaches with this recent perspective to flexible movement control. However, the separate quantification of the task-relevant information that is not shared or complementary between muscles (i.e. provided uniquely by individual muscles), is not possible using current analytical approaches. Intuitively, this task information encodes the functionally independent muscle activations that contribute uniquely to task performance, generated potentially by both central and peripheral sources ^18,20,21^. Although independent motor control mechanisms are well-established and can improve with training ^22,23^, the inclusion of this attribute in recent theoretical work marks a significant departure from traditional perspectives ^18^. Thus, here we will address this important current research gap by developing a methodology for the comprehensive quantification of unique task information in the muscle space.

A second consideration we address here concerns the coverage given by the muscle synergy concept to the contribution of whole muscle groups, rather than individual muscles, towards task-objectives. The motor redundancy problem motivating this concept describes how common neural inputs map to the task space through inter-muscular components ^13^. However, the end-effectors of this mapping operation are in fact individual muscles with their own unique anatomical attachments and activation timings ^16,24^, thus, as recognized by several lines of recent research ^9,15,18,25^, the muscle group may not be the smallest unit of modular control. The current muscle synergy concept therefore does not comprehensively describe the hierarchical structure of functionally cooperative muscles. Only more recently has the muscle synergy concept been formally applied at the intramuscular level ^25^, revealing that intramuscular synergies are independent of the muscles’ compartmentalized structure and may be complementary to inter-muscular analyses as a window into the neural control of movement ^26,27^. To incorporate this shift in perspective into current analytical approaches, here we aim to redefine the *‘working together’* notion of the muscle synergy to more comprehensively encapsulate this hierarchical characteristic of the human motor system.

In the current study, through a principled methodology, we aimed to probe the hierarchically structured functional architecture of the motor system, revealing salient features of movement modularity at multiple scales. To this end, we built upon traditional approaches and our recent innovations by directly including task parameters during muscle synergy extraction but here, by employing a Partial Information Decomposition (PID) (Fig.1A). With this proposed approach implemented in an established pipeline, we redefine the ‘*working together’* idea underpinning muscle synergies to characterize the hierarchical decomposition of task performance (*τ*) by functional modules (C1-C3 (Fig.1B)) that are comprised of diverse types of interactions (i.e. functionally -similar (redundant (pink intersection)), -complementary (synergistic (orange shading)) and -independent (unique (magenta and cyan intersections)) both between and within muscles (e.g. C2 and both C1 and C3 of Fig.1B respectively). Intuitively, synergistic information is the information shared by a given muscle pair ([*m*_*x*_, *m*_*y*_]) about *τ* that can only be gained by observing *m*_*x*_ and *m*_*y*_ together. On the other hand, redundant information is the task information shared by *m*_*x*_ and *m*_*y*_ that can be found in either alone. Finally, unique information is the task information provided by either *m*_*x*_ or *m*_*y*_ that is not found in the other. Hence, our framework can quantify any kind of functional relationship both inter- and intra-muscularly and align current muscle synergy analysis and its underlying *‘work together’* concept with the forefront of understanding on human movement modularity.

**Fig 1:**
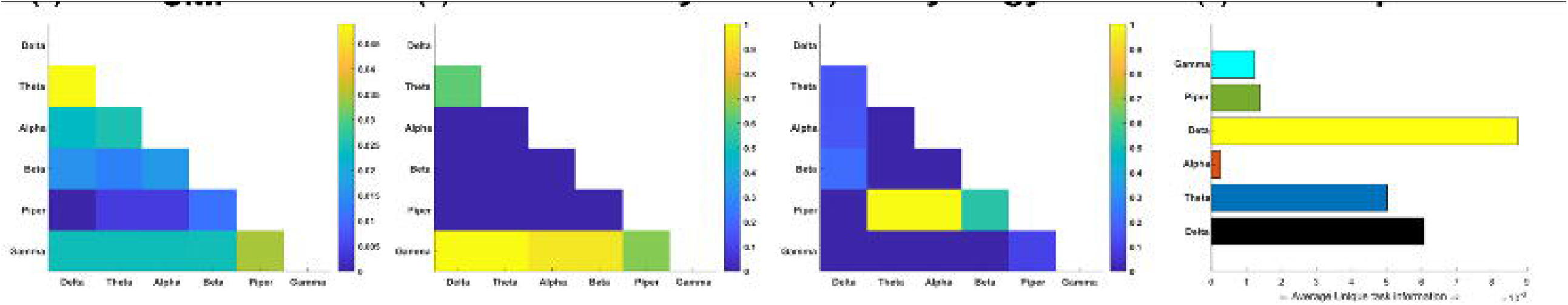
The proposed conceptual and methodological paradigm. **(A)** Top: Building on traditional muscle synergy analysis based on dimensionality reduction and a recent computational framework introducing an information-theoretic measure of net redundancy/synergy (pink-orange intersection) known as Co-Information ^28^, here we introduce Partial Information Decomposition (PID) ^29^, to more comprehensively quantify and functionally characterize muscle interactions. Bottom: The underlying premise of this framework builds on current approaches that quantify muscle covariations as shared variability (white-yellow intersection) and our previous work that dissects the task-relevant information from the task-irrelevant information (yellow intersection) and characterizes it as either functionally redundant (pink shading) or synergistic (orange shading) ^19^. By employing PID, here we incorporate the capability of separately quantifying redundant (pink intersection (**R**)), synergistic (orange shaded area (**S**)) and unique information (magenta (***U***_**1**_) and cyan (***U***_**2**_) intersections) from the shared information a given muscle pair ([**m**_**x**_, **m**_**y**_]) shares with a task parameter (**τ**) (see Equation 1.1-1.2 of Materials and methods section for further details). **(B)** To align our approach with recent theoretical work in the field ^18^, we redefine the *‘working together’* notion of muscles synergies to characterize a hierarchical decomposition of **τ** by functional modules comprised of diverse types of interactions both between and within muscles (e.g. C2 and both C1 and C3 respectively). **(C)** An overview of the proposed framework. (**a**) EMG data along with corresponding task parameters are recorded from human participants performing naturalistic movements. (**b**) PID is applied to all [m_x_, m_y_] and τ combinations, with the resulting information atoms then input into an established pipeline ^19,30^. (**c**) The output of this framework is low-dimensional components consisting of pairwise dependencies between muscles and their task- and participant-specific activations.

Towards the overall aim of this study, we applied the proposed framework to three human movement datasets (whole-body reaching – Dataset1, balancing on a balance board – Dataset2, and an object-lifting task – Dataset3) (Fig.1C), revealing generalizable patterns of diverse types of functional interactions both between and within muscles that encode distinct features of motor behavior. By redefining the muscle synergy idea as a hierarchical task decomposition implicitly in a computational approach, we provide crucial nuance and generalizability to motor control research to further biological insights and practical applications in the field. To supplement the work presented here, we have made available Matlab codes for readers to apply and simulate this framework: https://github.com/DelisLab/Muscle_PID.

## Results

### Principled mapping of muscle interactions to task performance

Our primary aim here is to probe the functional architecture of the motor system by establishing a principled method for quantifying task-redundant, -synergistic and unique informational dynamics underlying motor behavior, thus redefining the *‘working together’* concept of muscle synergies as a hierarchical task decomposition composed of diverse types of functional interaction both between and within muscles (Fig.1B). To achieve this, we firstly introduce our computational framework and apply it to pairs of EMG signals across time ([*m*_*x*_, *m*_*y*_]) and a corresponding, continuous task parameter (*τ*) (e.g. kinematics, dynamics etc.) (see ‘*Quantifying functionally diverse muscular interactions’* section in the Materials and methods) ^29^. The basic premise behind this approach is that the direct mapping of muscle interactions to task performance is crucial for understanding and principally quantifying their functional underpinnings. Unlike current approaches which first identify muscle couplings and then relate them to task performance, our proposed methodology firstly quantifies the task information carried by each muscle activation and then builds networks of common task information across muscles. The proposed Partial Information Decomposition (PID) approach parses the effects on *τ* by a pair of muscle activations ([*m*_*x*_, *m*_*y*_]) into four separate atoms of information (i.e. task-redundant (*R*), -synergistic (*S*) and two unique information components (*U*_*x*_ and *U*_*y*_)) by decomposing their joint mutual information (JMI) (see Fig.2(A) for a toy model). *R* is the predictive information between [*m*_*x*_, *m*_*y*_] about *τ* that can be found in either alone (e.g. *m*_*x*_ of Fig.2(A) redundant model alone fully predicts *τ*), encapsulating the portion of the [*m*_*x*_, *m*_*y*_] pairing that has a similar functional consequence. *S* on the other hand, is the predictive information provided by [*m*_*x*_, *m*_*y*_] about *τ* that is produced only when *m*_*x*_ and *m*_*y*_ are observed together (e.g. to predict *τ* in the synergistic system of Fig.2(A), both *m*_*x*_ and *m*_*y*_ need to be observed together), capturing the functional complementarity within the interaction. Finally, *U*_*x*_ and *U*_*y*_ is the predictive information within [*m*_*x*_, *m*_*y*_] about *τ* that is only present in *m*_*x*_ or *m*_*y*_ alone, capturing the unique contributions of the individual muscles within the functional interaction (e.g. *m*_*x*_ predicts a ‘L’ result in *τ* irrespective of the state of *m*_*y*_ and vice versa for predicting an ‘R’ outcome (Fig.2(A))). Thus, in contrast to the traditional conception of muscle synergies as co-variations across muscles, here instead we firstly extract shared task information between individual muscles, decompose each muscle interaction into different types of functional muscle covariation (see Fig.2(A)), and then determine common patterns of functional interactions across the muscle network (see Fig.3 for an overview of the framework).

**Fig 2:**
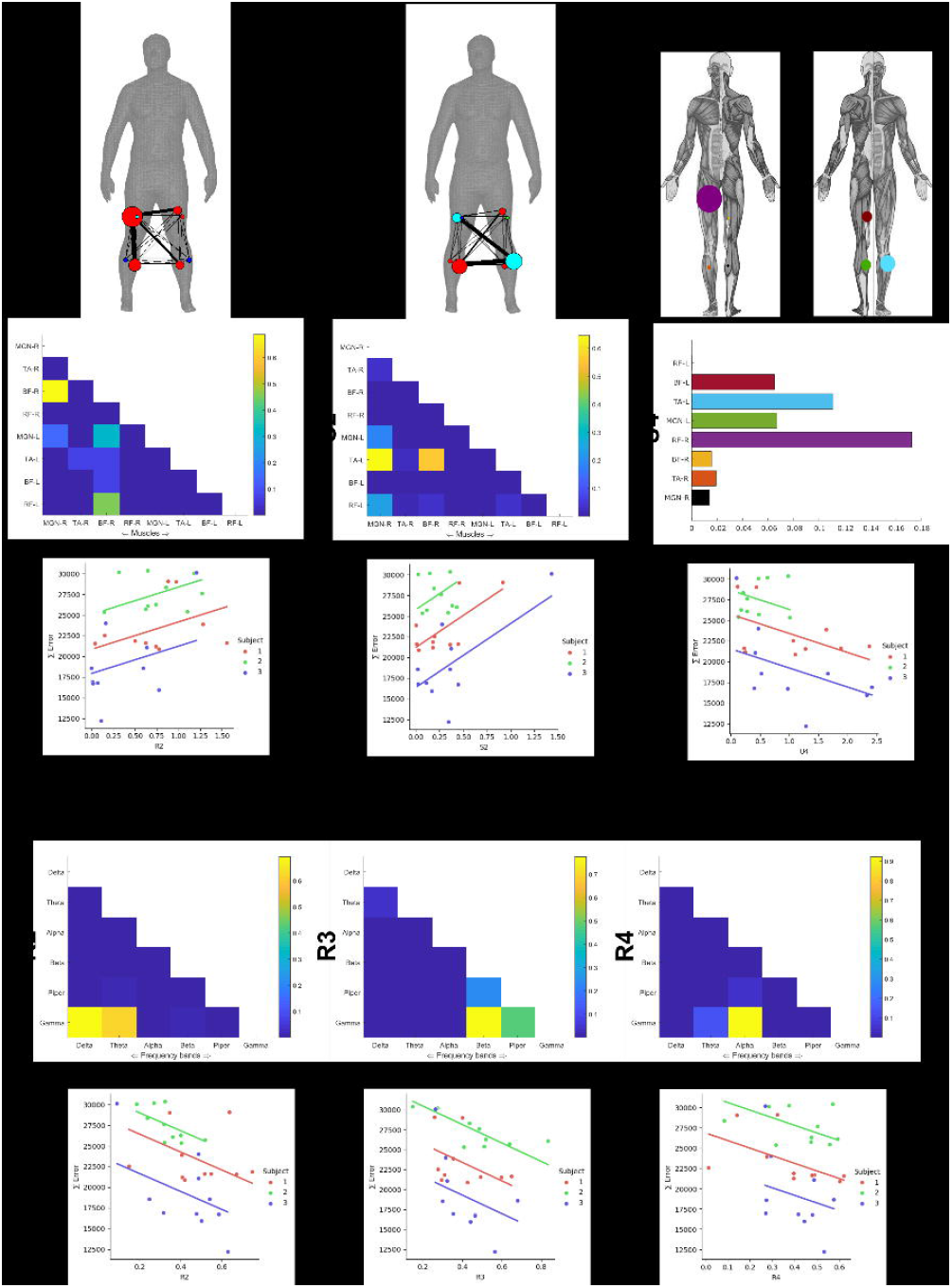
Toy simulation of different types of functional muscular interaction. **(A)** A toy model demonstrating how redundant, synergistic, and unique task information can be interpreted from the application of the PID approach to the muscle space. Four observations of a given muscle pair (*m*_*x*_ and *m*_*y*_) that can fall into two equiprobable on and off activation states and a corresponding task parameter (*τ*) describing left (L) or right (R) movement direction. Observing either *m*_*x*_ or *m*_*y*_ in the redundancy example gives 1 bit of information while observing both *m*_*x*_ and *m*_*y*_ together only in the synergy example gives 1 bit of information. Turning to the unique information example, when *m*_*y*_ is active in a specific way, it predicts a R outcome irrespective of the state of *m*_*x*_ and vice-versa. Thus, both *m*_*x*_ and *m*_*y*_ each provide task information that cannot be found in the other. **(B)** To further demonstrate the intuition behind our approach in recovering functional muscle relationships, we have carried out a toy simulation (see https://github.com/DelisLab/Muscle_PID) where we simulated two EMG signals (*M*_*x*_, *M*_*y*_) with a specified signal correlation (i.e. covariation in the average task-specific responses of the muscles) and noise correlation (i.e. covariations in the trial-to-trial responses of the muscles). The joint responses of *M*_*x*_ and *M*_*y*_ are plotted for different combinations of positive, negative and null signal and noise correlation where the ellipses illustrate the direction and overlap of the muscles responses. **(C)** PID was applied to *M*_*x*_ and *M*_*y*_ at a range of positive and negative signal and noise correlations, describing how redundancy, synergy and unique information vary with respect to these task encoding mechanisms. The PID values presented are normalized with respect to the JMI, thus illustrating the proportional contributions of each interaction type. For unique information, the average over *M*_*x*_ and *M*_*y*_ in each instance is displayed.

**Fig 3:**
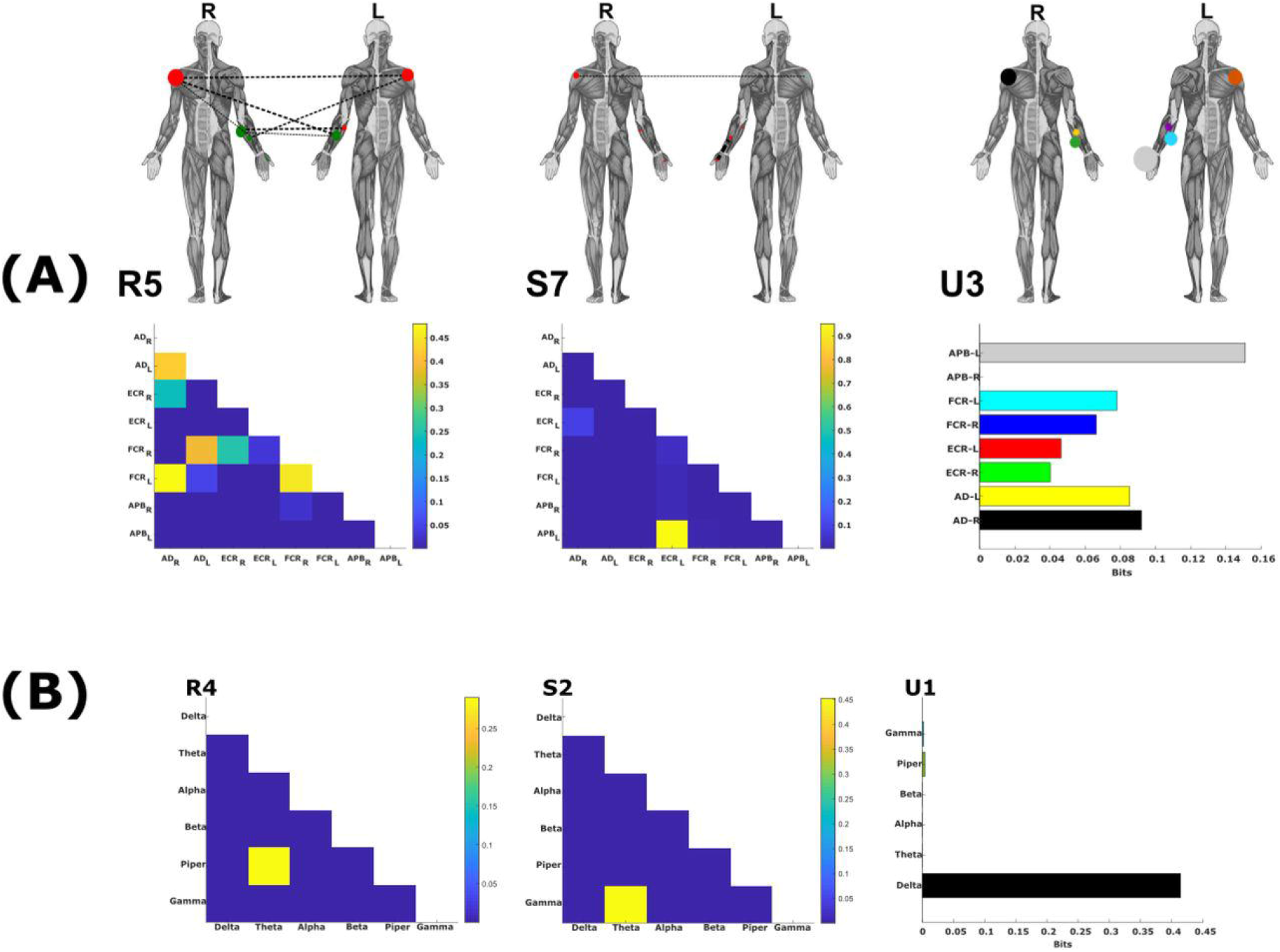
A summarized overview of the NIF pipeline. **(A)** EMG data is captured from human participants performing naturalistic movements. **(B)** The joint mutual information (JMI) between all muscle pair and continuous task parameter combinations is decomposed using the partial information decomposition framework, resulting in separate networks of redundant (R), synergistic (S), and unique muscle interactions (U1, U2). **(C)** Each network is sparsified with respect to its percolation threshold, identifying statistically significant functional connections. **(D)** A hierarchical community detection essentially unravels and identifies overlapping clusters of network dependencies. **(E)** The optimal number of clusters identified serves as the input parameter into dimensionality reduction, where like current approaches, low-dimensional components of muscular interactions along with task- and participant-specific activations are extracted.

### Simulating functionally diverse muscle interactions reveal close associations with neural coding concepts

To provide further intuition on the use case of this approach in recovering functional relationships between muscles, we also implemented a toy simulation in Matlab (see https://github.com/DelisLab/Muscle_PID) (Fig.2(B-C)). To briefly summarise, we simulated multiple trials (N=100) of two EMG signals (*M*_*x*_, *M*_*y*_) by summing a combination of sinusoids of frequencies commonly found in EMGs (i.e. 20-150Hz). Additionally, we included a shared encoding between the EMGs for a simulated binary task parameter and then adjusted their relative tuning towards this task parameter such that their average task-specific responses could have positive (similar), negative (complementary) or null (independent) correlations (i.e. signal correlation). Further, we then injected separate Gaussian noise signals into both EMGs with a specified level of trial-to-trial covariation (i.e. noise correlation).

Fig.2(B) illustrates the average responses of *M*_*x*_ and *M*_*y*_ with respect to one another for different signs of signal and noise correlation. The ellipses on each plot here indicate the distribution of each muscles’ average responses, together illustrating their relative tuning direction and overlap.

To determine how these patterns of joint response are reflected within our framework, we applied PID to *M*_*x*_ and *M*_*y*_ at various ranges of signal and noise correlation with respect to the corresponding task parameter (Fig.2(C)). Corresponding well with our previous insights here (Fig.2(B)), we found that when signal and noise correlations have the same sign (see top right and bottom left panels in Fig.2(B)), the information provided by the muscle activation relationship is redundant. In this case, e.g. a high value of *M*_*x*_ provides common task information with e.g. a low value of *M*_*y*_. On the contrary, when signal and noise correlations have the opposite sign (see top left and bottom right panels in Fig.2(B)), the information provided by the activation relationship is synergistic. In this case, it is the relationship between the two muscles that increases the task information. For example, knowing that the activation of *M*_*x*_ is high while the activation of *M*_*y*_ is low provides more task information (and consequently better task discrimination) than observing the two muscle activations independently. Finally, when signal (or noise) correlations are close to zero (see e.g. middle panel in Fig.2(B)), task information is conveyed uniquely by each muscle and not their relationship. In this example (middle panel), only the activation of *M*_*x*_ conveys information about the task performed.

Altogether, the results of this toy simulation demonstrate how our approach can capture functional muscle relationships of any kind in a way that aligns closely with established research on the mechanisms of neural coding ^31–33^.

### The Network-Information Framework pipeline

Continuing, following this decomposition, we then ran the separate PID atoms through an established pipeline ^19,30^, referred to as the Network-Information framework (NIF) (Fig.3(A-E)) (see Materials and methods for detailed breakdown). The purpose of the NIF is to produce functionally and physiologically relevant and interpretable low-dimensional components of muscular interactions underlying coordinated movements. The following briefly summarizes the main steps of the pipeline:

1. To produce a comprehensive network of functional muscle interactions, we applied the PID framework over all unique [*m*_*x*_, *m*_*y*_] and *τ* combinations for each participant (Fig.3(B)). This iterative procedure results in a multiplex network, with each layer consisting of all functional dependencies between muscle pairs for a particular PID atom, *τ* and participant.
2. To determine the statistically significant interactions at the network level, we applied a modified percolation analysis to each layer of the multiplex network (Fig.3(C)) ^6^.
3. To determine the optimal number of clusters to extract from structurally nested networks, we employed a link-based community detection protocol based on a modularity maximization cost-function (Fig.3(D)) ^34–36^.
4. The optimal cluster count was then used as the input parameter into dimensionality reduction, namely a projective non-negative matrix factorization (PNMF) algorithm ^37^, to extract patterns of muscle connectivity along with their task- and participant-specific activations (Fig.3(E)).

### Hierarchical and functionally diverse muscular interactions underly motor behavior

To demonstrate the proposed PID approach, we present an example output from an application to the EMG recordings of a single participant (across all trials) from Dataset 1 with respect to the combination of the reaching finger kinematic coordinates (i.e. X*Y*Z). This participant was instructed to perform a total of 72 different randomly selected whole-body point-to-point reaching tasks for ∼2160 trials (see Fig.4(a)) and *‘Data acquisition and experimental conditions’* section of the Materials and methods section). In Fig.5, we illustrate the redundant (*R* – 5b), synergistic (*S* - 5c), and unique (*U*_*x*_ and *U*_*y*_ - 5d) interactions between muscles, as well as their sum total which they are all normalized by, (i.e. JMI −5a). Human body models accompanying each of the JMI, *R* and *S* muscle networks illustrate the strongest interactions between muscles (indicated by edge-width) ^40^, the muscle subnetworks (node color) and the network centrality (a measure of a muscle’s relative importance indicated by node size) (see ‘*Subnetwork analysis’* section of Materials and methods) ^34,41,42^. The *U*_*x*_ and *U*_*y*_ terms are not considered as a muscle coupling, as they encode the task information present in one muscle that is not present in another and vice versa, and so instead the average unique information (*U*_*xy*_) for each muscle is presented as a bar graph that is color coded instead to illustrate specific bodily regions.

**Fig 4:**
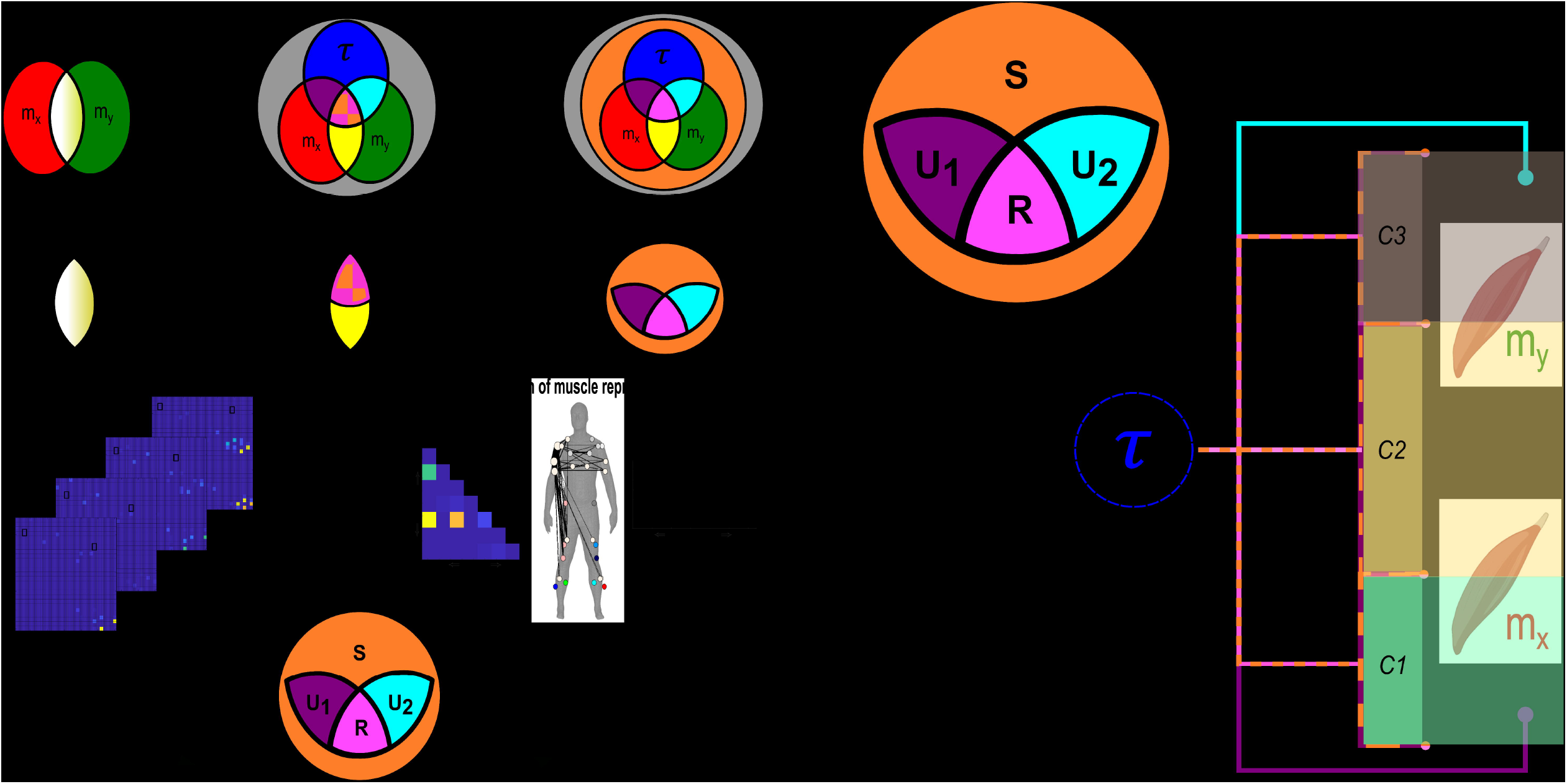
The datasets used in the application of the proposed framework in the current study. **(a) Dataset 1:** Healthy adult participants performed whole-body reaching tasks in various heights and directions while EMG and kinematics were captured across the body ^38^. **(b) Dataset 2:** Healthy adult participants performed 10 trials of balancing on a balance board while EMG was captured among the bilateral lower-limb flexors and extensors simultaneously to the horizontal angular displacement of the balance board. **(c) Dataset 3:** Healthy younger and older adults performed a reach-grasp-lift-hold and replace task of both light and heavy objects while EMG from the arm musculature bilaterally were captured along with load and grip forces on the grasped object ^39^. For full details on the experimental setup of these datasets, see *‘Data acquisition and experimental conditions’* section of the Materials and methods.

**Fig 5:**
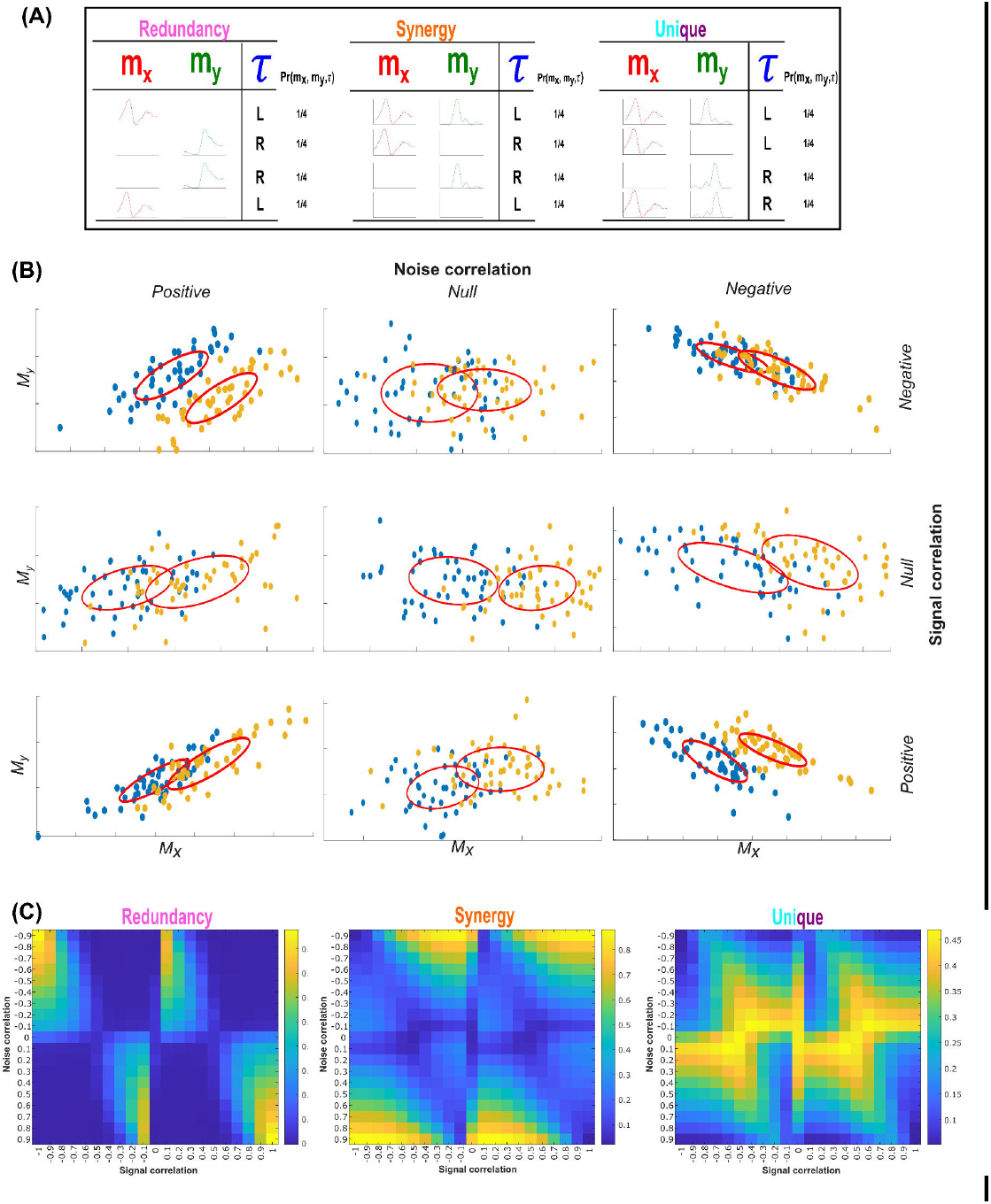
An example application of the proposed framework. Application of the proposed framework to the intermuscular space of a single participant performing multiple trials of various whole-body reaching tasks (Dataset 1). The Joint mutual information (JMI) **(a)** and its informational components, redundant **(b)**, synergistic **(c)** and unique information **(d)** were determined with respect to the combination of the reaching finger kinematic XYZ coordinates (i.e. X*Y*Z). Values for *R, S* and *U*_*xy*_ were normalised with respect to the presented JMI values ^43^. Human body models accompanying each representation in **(a)**-**(c)** illustrate the strongest connectivities (edge-width) ^40^, the subnetwork community structure (node color) and network centrality (relative node size) ^34,41,42^. The *U*_*x*_ and *U*_*y*_ terms are not considered as a muscle coupling, as they encode the task information present in one muscle that is not present in another and vice versa, and so instead the average unique information (*U*_*xy*_) for each muscle (LLB=purple, LUB=green, RLB=red, RUB=black) is presented as a bar graph. The accompanying human body model to the left illustrates the predominant muscles (color coded to represent their bodily region) that encoded unique information about the right, reaching finger kinematic.

The JMI network is comprised of a single submodule (red nodes) with the greatest network centrality among the reaching-side biceps brachii and lateral triceps (Fig.5(a)). These prime-mover muscles are central as basically all other muscles are functionally coupled to them, while the contralateral biceps and triceps mirror this organization to some extent with further interactions with the left biceps femoris and bilateral rectus femoris. When these dependencies are decomposed into their PID components (Fig.5(b-d)), we reveal a more complex functional architecture underlying whole-body reaching movements. The functionally -similar (redundant) and -complementary (synergistic) networks consist of multiple overlapping subnetworks widespread across the body (Fig.5(b-c)). Amongst the redundant subnetworks, several muscles demonstrate a high level of centrality (e.g. right pectoralis, trapezius, biceps femoris and vastus lateralis, left tibial musculature and bilateral gluteus maximus)(Fig.5(b)), while synergistic subnetworks demonstrate a more even spread of functional importance (Fig.5(c)). The synergistic network consisted of functional muscle couplings that were essentially counterfactual to the connectivity of the redundant network (i.e. all connections not present within the redundant network were present within the synergistic network and vice versa), thus illustrating the crucial cross-module connectivities that functionally integrate complementary information across the muscle network. Interestingly, the redundant prime-movers in Fig.5.(b) did not contain much unique task information (Fig.5(d)), while muscles displaying considerable functional independence did not feature prominently in the redundant networks (e.g. right triceps brachii, anterior and posterior deltoid). This suggests that although different types of interaction co-occur between muscles, their proportional contributions map strongly to the muscles’ physiological function in the context of the task demands (i.e. reaching-side shoulder musculature require more selective control to guide the arm to specific targets (see Fig.4(a)).

Next, to elucidate the functional interactions within muscles underlying whole-body reaching movements, we applied the proposed methodology to pairwise combinations of amplitude signals from six frequency-bands ([*f*_*x*_, *f*_*y*_]) (Delta [0.1-4 Hz], Theta [4-8 Hz], Alpha [8-12 Hz], Beta [12-30 Hz], Low-Gamma (Piper rhythm) [30-60 Hz], High Gamma (Gamma) [60-80 Hz]) (see ‘*Quantifying diverse muscular interactions in the task space’* and ‘*Data pre-processing’* sections of the Materials and methods) extracted from the right anterior deltoid muscle. This computation serves as a nonlinear measure of coherence decomposed into its task-relevant informational constituents, which we used here to determine the multifarious effects of [*f*_*x*_, *f*_*y*_] on the right, reaching finger anteroposterior kinematic (Fig.6). The intramuscular JMI network comprised of a mixture of dependencies between distinct oscillations (Fig.6(a)), most prominently between the Delta-Theta rhythms. When we decomposed their shared task information, the rhythmic activities of the right anterior deltoid presented a more distinguishable encoding of task performance. Gamma amplitudes were functionally similar in their encoding of the finger kinematic with respect to all other frequency bands while a separate Delta-Theta coupling was also had functionally similar consequences (Fig.6(b)). The Piper rhythm comprised of task-synergistic information in the anterior deltoid muscle when coupled with the Theta and Alpha oscillations (Fig.6(c)). Meanwhile, the amplitude of beta oscillations provided the most functionally independent information about the reaching finger kinematic on average, followed by Delta and Theta amplitudes (Fig.6(d)).

**Fig 6:**
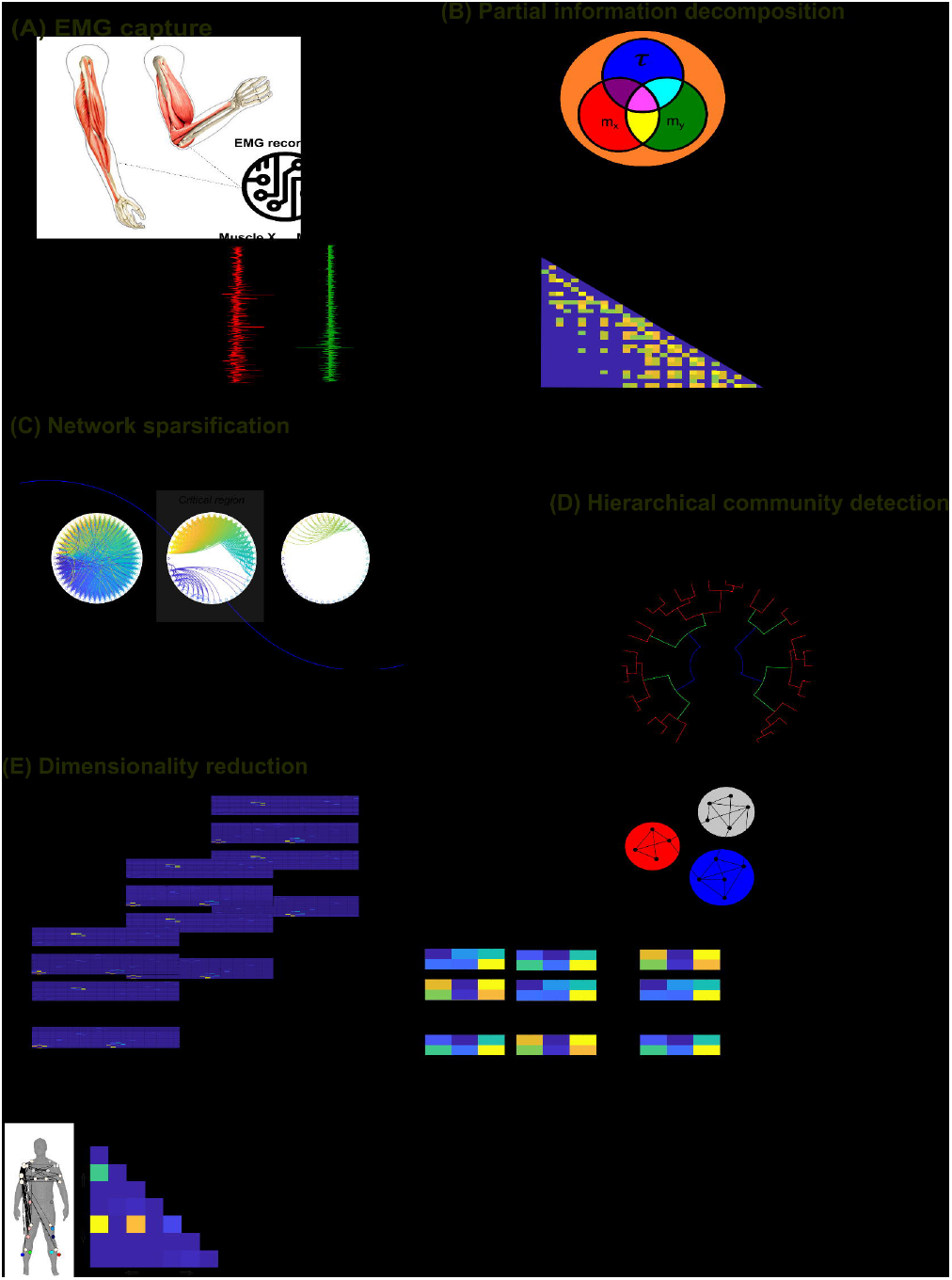
Application of the proposed framework to frequency-specific amplitudes in the reaching side anterior deltoid muscle of a single participant performing multiple trials of various whole-body reaching tasks (Dataset1). The Joint mutual information (JMI) **(a)** and its informational components, redundant **(b)**, synergistic **(c)** and unique information **(d)** were determined with respect to the reaching finger anterior-posterior kinematic coordinate. Values for *R, S* and *U*_*xy*_ were normalised with respect to the presented JMI values ^43^. The *U*_*x*_ and *U*_*y*_ terms are not considered as a coupling, as they encode the task information present in one oscillation that is not present in another and vice versa, and so instead the average unique information (*U*_*xy*_) for each frequency-band are presented together as a bar graph.

These finding provide strong evidence towards the basic premise of and motivation for this framework in redefining the muscle synergy concept as a hierarchical decomposition of motor behavior by functionally diverse muscle interactions, evidence that links well with emerging findings in the field [14, 15].

### Generalizable components of functionally diverse inter- and intra-muscular interactions

Having demonstrated the presence of the diverse types of functional interactions both between and within muscles, we then sought to extract motor components that are generalizable beyond any individual participant and task.

Beginning with intermuscular components, we identified four *R* and *S* (R1-R4 and S1-S4) and three *U*_*xy*_ (U1-U3) components with respect to the XYZ (anterior-posterior, medio-lateral vertical directions) coordinates of 21 kinematic markers (21 x 3 dimensions = 63 task parameters in total) across the body of three participants performing whole-body reaching movements (>2000 trials each). For brevity here, we provide illustrations of the output in the supplementary materials (Supp. Fig.1-3). To examine the generalizability of these components, in a leave-one-out cross validation procedure, we removed an individual task parameter or participant from the input data and then re-extracted the same number of components and computed correlations between this data subsets’ output and the full dataset’s output (see *‘Examining the generalizability of the extracted components’* section of the Materials and methods). We found an almost perfect concordance between the presented intermuscular components and those extracted from a subset of the data (∼0.99 average correlation). The robustness of these components exceeds previous implementations of the NIF ^19,30^, where a high level of concordance was also found.

Turning to the intramuscular space, applying the proposed approach within all 30 muscles with respect to 63 kinematic coordinates from the three participants of Dataset 1 revealed four *R* (R1-R4 (Supp. fig.4)) and three *S* (S1-S3, (Supp. Fig.5)) and *U*_*xy*_ (U1-U3 (Supp. fig.6)) components. The generalizability of these intramuscular components was proven at the intramuscular level with ∼0.99 correlation typically for both individual tasks and participants among *R* and *S* networks. A slightly lower concordance among *U*_*xy*_ (r=0.9) was found when an individual participant was removed from the input data.

### Hierarchies of functional muscle interactions encode distinct motor features

Finally, having quantified diverse types of functional muscle interaction at both inter- and intra-muscular scales, we then investigated the functional relevance of the extracted muscle networks at each of these scales. Specifically, we asked if the identified muscle interactions are reflective of motor performance (balance – Dataset2) and motor decline with age (object lifting – Dataset3).

#### Functionally diverse intermuscular interactions reflect balance performance

To begin with dataset 2, we identified and extracted five *R*, four *S* and four *U*_*xy*_ intermuscular components. We then used the trial-specific activations from the extracted components for each participant (normalized with respect to their corresponding JMI) to predict motor performance in each trial (i.e. the total balance board error calculated as the sum of absolute deviations from 0 degrees on the horizontal plane of the balance board) (∑Error) (see ‘*Salient features of motor performance’* of the Materials and methods section). As multiple observations for the same participant were present within these vectors (n=10 trials each), we determined associations using repeated measures correlation ^44^, a measure of linear correlation that models participant-specific clustering in the data.

We found two intermuscular interactions (S2 with synergistic couplings between both BF-R and MGN-R, and the left tibialis anterior (TA-L), r=0.64, p=0.0003 and R2 with redundant couplings between the right medial-gastrocnemius (MGN-R) and biceps femoris (BF-R) and BF-R and left rectus femoris (RF-L), r=0.64, p=0.0003) (Fig.7(A)). Interestingly, muscle couplings representing redundant and synergistic interactions here did not appear prominently in the motor component U4 contributing uniquely to task performance. In contrast, the unique task information in the right rectus femoris (RF-R) and to a lesser extent TA-L (together possibly representing their important roles in crossed-extensor reflex actions) was related to a significant reduction in ∑Error (r=-0.44, p= 0.02), i.e. predicting better balance performance (Fig.7(A)).

**Fig 7:**
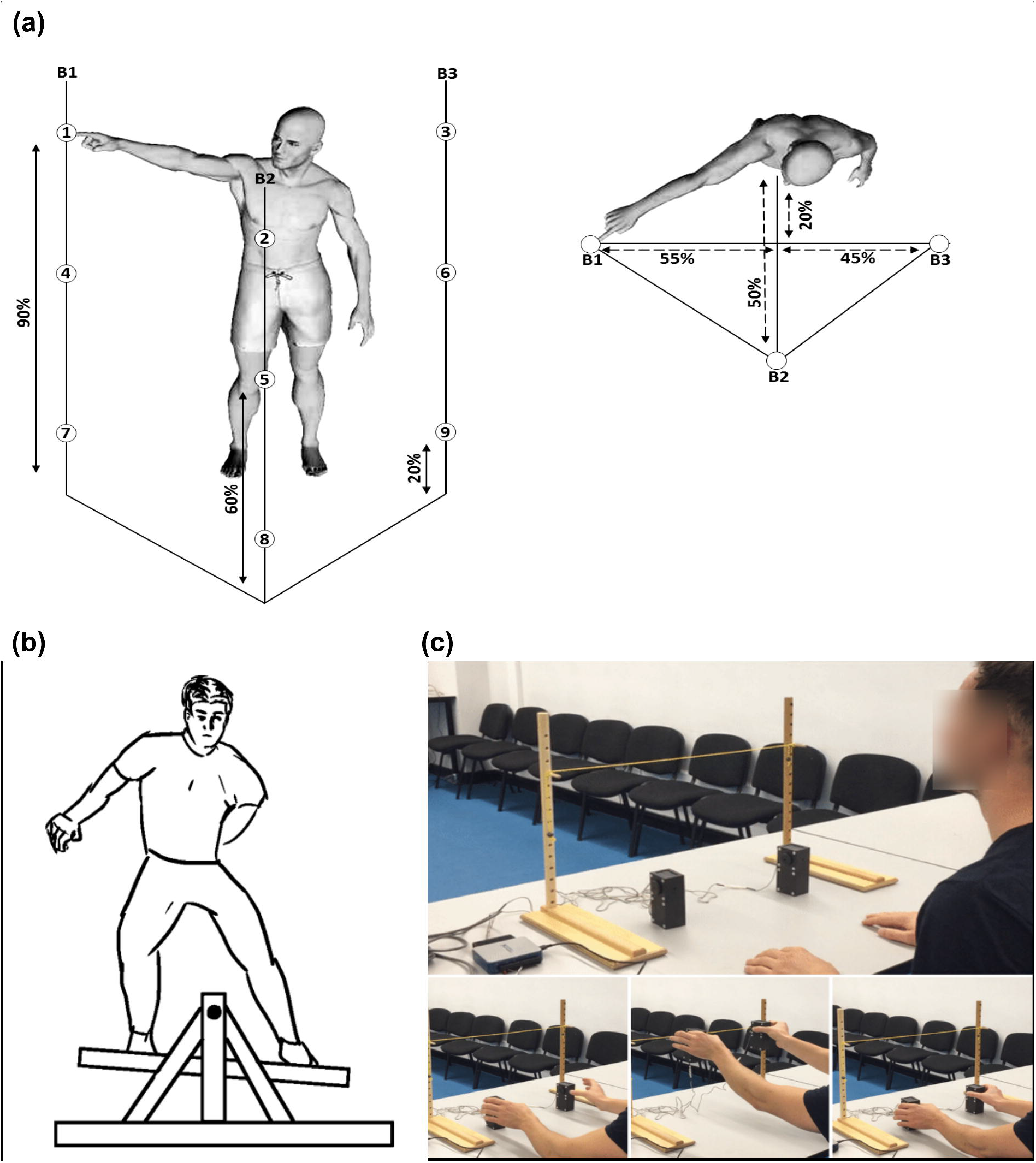
Application of the proposed framework to multiple trials on a balance board. **(A)** The intermuscular components (R2, S2 and U4) determined to have a significant (P<0.05) correlation with balance performance (∑Error) across trials in Dataset2. Top: each adjacency matrix, the human body models illustrate the network structure with relative edge thickness, node size and color reflecting connection strength, importance and sub-modularity respectively ^34,41,42^. Bottom: Scatterplots illustrating the corresponding repeated measures correlation with balance performance. **(B)** Top: The intramuscular components (R2, R3 and R4) determined to have a significant (P<0.05) correlation with balance performance (∑Error) across trials in dataset 2. Bottom: Scatterplot of the corresponding repeated measures correlation outputs.

#### Functionally similar intramuscular interactions reflect balance performance

Turning to the intramuscular space, we identified and extracted five *R* and three *S* and *U*_*xy*_ from Dataset2 and found three of these components significantly correlated with ∑Error across trials (see ‘*Salient features of motor performance’* of the Materials and methods section) (Fig.7(B)). These components all represented redundant amplitude couplings (R2-R4), consisting exclusively of specific coherences between the gamma band and other lower frequency amplitudes (delta and theta in R2, beta and piper in R3 and, alpha in R4), and were negatively correlated with ∑Error (i.e. increased involvement of intramuscular modules resulted in improved balance performance, r=-0.45, p=0.015 for R2, r=-0.51, p=0.006 for R3, and r=0.42, p=0.03 for R4).

#### The activation variability of a combination of bilateral muscle networks predicts age differences in bimanual object-lifting

Next, we asked which muscle interactions may underpin differences in motor performance resulting from ageing. In the motor control literature, older adults have been shown to exhibit greater motor variability compared to young cohorts ^45^, leading to behavioral inconsistency. To answer this question, we applied our proposed approach to Dataset 3 ^39^, consisting of EMG recordings (8 upper-limb muscles bilaterally) from healthy young (N=14) and older (N=18) participants performing a bimanual reach-grasp-hold task of both light and heavy objects (see Fig.4(c) and ‘*Data acquisition and experimental conditions’* section of the Materials and methods).

Applying PID to all pairs of EMG signals to predict the bilateral grip and load forces (i.e. 4 task parameters) the participants exerted, we identified and extracted five *R*, seven *S* and three *U*_*xy*_ intermuscular components. To investigate how this variability would manifest in the extracted functional muscle patterns, we defined a measure of motor variability in the activation of muscle or frequency couplings (∑Error) and applied it to the extracted components (see *‘Salient features of motor performance’* of the Materials and methods section). These vectors were then employed as the predictors in a binary logistic regression model against participants’ age group (Young=0 vs. Old=1). This procedure indicated that a combination of the 5^th^ *R* (β= −2.12±0.801, p<0.01), 7^th^ *S* (β= 1.36±0.704, p=0.053) and 3^rd^ *U*_*xy*_ (β= 0.964±0.445, p<0.05) motor components were optimal in predicting age group (Fig.8(A)), classifying 75% of participants correctly.

**Fig 8:**
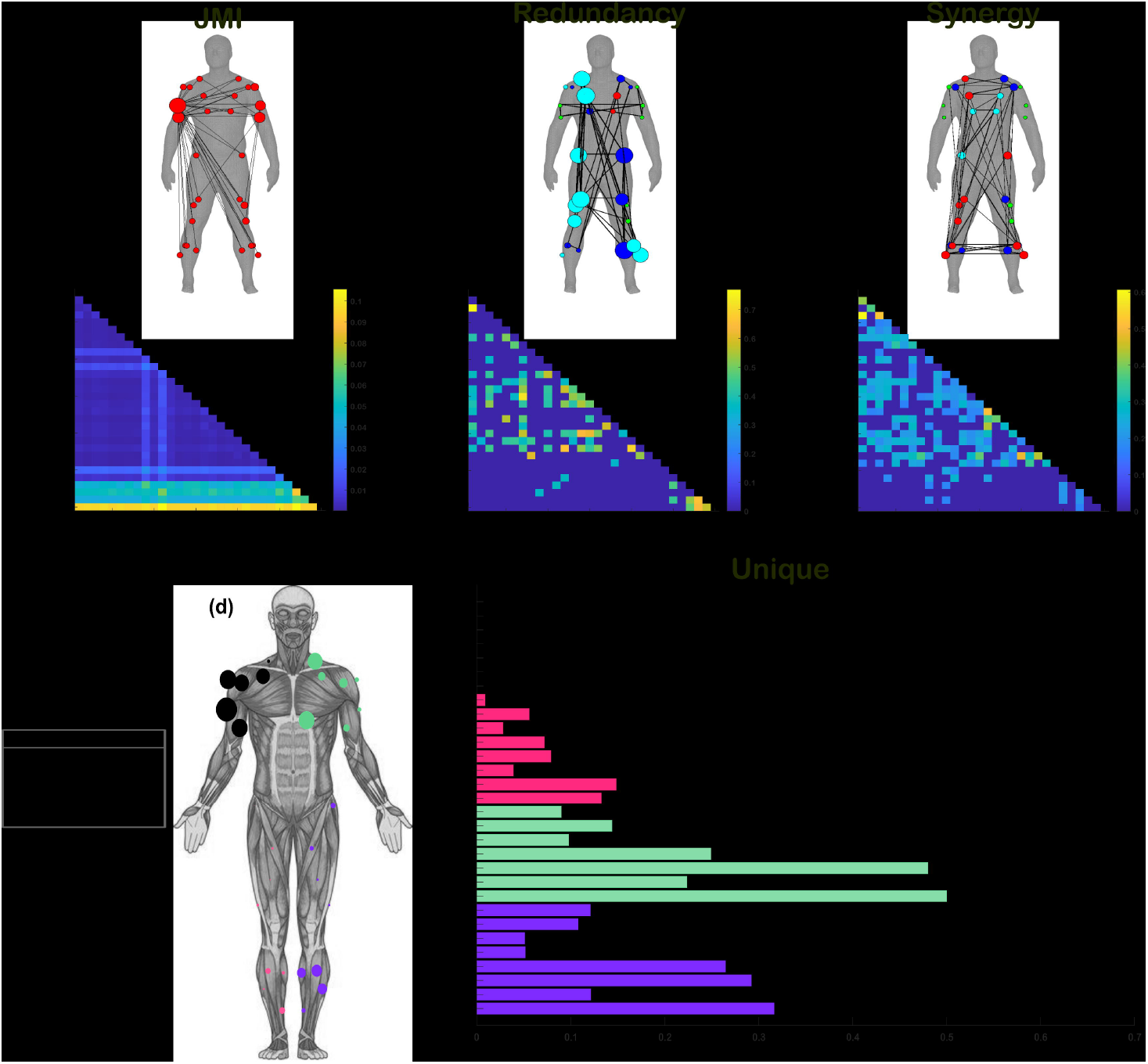
Application of the proposed framework to young and older adults performing reach-and-grasp tasks. **(A)** The intermuscular components (R5, S7, U3) whose underlying recruitment variability (∑ *Error*) was significantly predictive of participants’ age group. All three components formed part of a binary logistic regression model (R5: [β= −2.12±0.801, p<0.01], S7: [β= 1.36±0.704, p=0.053], U3: [β= 0.964±0.445, p<0.05]) and classified 75% of participants correctly. Above the adjacency matrices, human body models illustrate the network connectivity with relative edge width, node size and color representing the connection strength, muscle importance and sub-modularity ^34,41,42^. **(B)** The intramuscular components (R4, S2, U1) whose underlying recruitment variability (∑ *Error*) was significantly associated with participants’ age group. All three components formed part of a binary logistic regression model (R4: [β= −0.557±0.313, p=0.075], S2: [β= −11.75±4.6, p=0.011], U1: [β= 1.94±0.982, p<0.05]) and classified 84.4% of participants correctly.

#### The activation variability of a combination of functionally diverse intramuscular rhythms predict age differences in bimanual object-lifting

When applied in the same way to the corresponding intramuscular level of Dataset 3, we identified and extracted five *R*, four *S* and four *U*_*xy*_ rhythmic components. Participants’ age group was optimally predicted by a combination of the 4^th^ *R* (β= −0.557±0.313, p=0.075), the 2^nd^ *S* (β= −11.75±4.6, p=0.011) and the 1^st^ *U*_*xy*_ (β= 1.94±0.982, p<0.05), which classified 84.4% of participants correctly (Fig.8(B)). Taken together, these findings suggest that concomitant increases and decreases in the variability of functionally diverse muscle interactions across scales characterize aging-induced changes in upper limb motor function.

## Discussion

In this study, we aimed to probe a recently proposed functional architecture for the human motor system by redefining the muscle synergy concept. To this end, we proposed a computational approach to muscle synergy analysis that separately and simultaneously quantifies the redundant, synergistic, and unique contributions of muscle interactions to task performance at both intermuscular (i.e. pairs of muscle activations) and intramuscular (i.e. pairs of rhythmic activities) levels. This approach presents a nuanced perspective to the *‘work together’* idea of muscle synergies, conceptualizing it instead as the hierarchical decomposition of motor behavior by functional modules comprised of diverse types of interactions both between and within muscles. It also builds on a previously established pipeline for the extraction of task-relevant muscle interaction patterns whose basic premise is that muscle interactions should be directly mapped to task performance to understand their specific functional underpinnings. Our innovation here goes further by enabling the quantification of the independent contributions of muscles (or intramuscular oscillations) to task performance, thus integrating recent theoretical perspectives on motor modularity into muscle synergy analysis. In firstly applying this framework in a toy simulation, we found that our approach is capable of recovering functional muscle relationships in an intuitive way. From applying our framework to benchmark datasets, we then demonstrated how the decomposition of task-relevant muscle combinations reveals complex functional architectures underlying everyday human movements. These functional architectures displayed a highly nested network structure of functionally similar and complementary information processing between muscles along with their independent functional contributions (as depicted in fig.1.2). We then investigated the functional interactions at the intramuscular level between frequency-specific oscillatory amplitudes, revealing that functional modularity is a scale-invariant characteristic fundamental to movement construction. We found that the extracted inter- and intra-muscular components were consistently generalizable beyond any data subset and across disparate motor tasks such as whole-body reaching and balancing tasks. Finally, we showed that these motor components were differentially correlated with salient motor features including motor adaptation and age group, suggesting they offer complementary windows into movement control. Thus, we present the proposed framework as a powerful tool for fundamental mechanistic investigations from a the neural coding perspective on human action. As demonstrated here, this analytical tool can be used to a) characterize the functional outcomes of muscle interactions and compare them across populations or conditions and, b) identify physiological markers of motor performance, learning or impairment.

The proposed methodology aligns with recent theoretical innovations in the field proposing functional modularity both within- and between muscles as a mechanism for simplified and flexible movement control ^18^. Within this theoretical framework, motoneurons affiliated with overlapping functional groups may innervate the same or different muscles, while individual muscles can also be independently controlled concomitantly. This proposition implies a diverse range of hierarchical muscular interactions including muscles working together towards functionally similar (redundant), complementary (synergistic) and independent (unique) task-goals ^14,15,17^. Inspired by the neural coding literature ^31–33^, by posing the degrees-of-freedom problem of coordinated movement as the problem of information sharing across muscle networks, our framework effectively reveals this hierarchical and diverse functional architecture. To elaborate, the extracted motor components captured multiple, co-occurring subnetworks of muscles (or intramuscular rhythms) forming overlapping functional groups. The functional nestedness unveiled here suggests that the motor system generates hierarchical task representations whose dimensions are folded into one another to simplify movement control. In effect, and as conceptualized here as *‘working together’*, this suggests that the motor system adaptively decomposes the problem of task execution into manageable sub-tasks addressed through hierarchical recursion, interestingly mirroring dynamic programming architectures ^46^. Continuing, several of these patterns were shared across interaction types (e.g. Supp. Fig. 1-3), indicating diverse information processing indeed occurred concomitantly within the same muscle networks. Many of these motor components were centrally defined or were more widespread across the body. These networks of functional muscle connectivity across the body are underpinned by polysynaptic communication pathways involving feedforward and feedback processes ^21^. Among these communication pathways, both the centralized and peripherally-sourced routing of task information globally integrate muscle functionality across the body while diffusive processes prevail more locally, promoting a segregation of functional roles ^20,47–49^. Moreover, the unique adherence of top-down and bottom-up motor processes to specific task features enhances the selectivity of sensorimotor representations and consequently the adaptability of movement control ^21,50^. The results of our toy simulation suggest that the capacity for muscles to be uniquely adherent to specific task-objectives however are constrained to specific ranges of noise correlation (see Fig.2(C)), suggesting limits to the independent control of muscles ^51^. Future work incorporating network communication models with the capacity to integrate these mechanisms of information transfer may be fruitful for understanding how motor signals are broadcast throughout the nervous system ^49^.

The separate quantification of unique, redundant, and synergistic task information proposed here brought about interesting insights into human motor control. For instance, we were able to identify the functional underpinnings of increased motor variability in older adults. We found variability in functionally complementary intermuscular couplings increased with age but decreased intramuscularly with age also (Fig.8), suggesting ageing has differential effects on functionally integrative mechanisms across scales. Meanwhile, variability in functionally redundant and independent networks reduced and increased respectively across both scales in the older adults group. These findings firstly contest the notion of increased motor variability with aging being simply a manifestation of motor noise, but demonstrate in fact that this variability manifests in functionally relevant channels of muscle interaction. These insights also support empirical work highlighting functional integration as a compensatory mechanism in individuals with age-related neurodegeneration ^52^, going further in showing that this supplementary integration is not present across all scales but, in fact, may come at the expense of other scales. Finally, the fact that all three interaction types each had unique explanatory information about ageing highlights the comprehensive characterization of motor control our approach can provide about multifaceted health conditions. Future work identifying the exact neural underpinnings of these distinct types of muscle interaction will further bolster the clinical insights of this approach. Continuing, we were also able to show that diverse types of task information are not isolated to specific muscle couplings or rhythms but are dynamically generated by various oscillatory signals to meet task demands. Among intramuscular *R*, gamma amplitudes were repeatedly associated with improved balance performance across participants when coupled in a functionally similar way with alpha- and beta-bands (Fig.7(B)). The alpha and beta-bands have been the subject of focus in several studies showing their crucial role in muscle synergy generation and monitoring ^53–55^. Here we add to this line of research by uncovering a potentially crucial role of higher gamma oscillations in movement control, showing that gamma amplitudes alone can provide much of the functionally relevant information provided by these lower-frequency bands. In sum, the application of our approach brought about meaningful insights into human motor control.

Despite the strong base of evidence demonstrating the capacity of the individual muscle to augment whole movement patterns ^20,56,57^, muscle synergy research has generally focused on muscle clustering’s. Our nuanced definition and subsequent findings here support recent formal applications of the muscle synergy concept to the intramuscular level, where task-specific modules have been identified ^25–27^. Of note, past research suggested that intramuscular modules represent spinal-level circuitry while the intermuscular space primarily captures supraspinal mechanisms ^27^. Indeed, this observation is supported by work from several research groups using different analytical techniques ^58–60^. Here, we consistently found that the intramuscular level represented aspects of motor behavior indicative of superior task performance and proficiency (Fig.7-8), while the intermuscular level mainly represented decrements in performance and compensatory mechanisms (Fig.7-8). This opposing pattern intuitively aligns with this recent work, as erroneous performance requires more frequent intervention by supraspinal mechanisms while more effective movement can be coordinated automatously by spinal circuitry. An exception to this pattern however lies in the independent control of muscles quantified here as unique task information which demonstrated a correlation with better task performance inter-muscularly (Fig.7(A)) and of which increased fluctuations were consistently related to older age across both scales (Fig.8). This independent control mechanism is likely invoked where coarse-grained control mechanisms are insufficient, allowing for the necessary selectivity to maintain task performance. This suggests that variability in independent control mechanisms both between and within muscles plays a major role in the manifestation of (and/or compensation for) behavioral inconsistency among older adults. However, as other related work also proposed ^16,17,54,58^, our findings here (Fig.7-8) suggest that there is not a clear distinction between the neural substrates underlying inter- and intra-muscular dynamics, and that they likely reflect the contributions of multiple neural substrates simultaneously albeit to different extents, together holistically representing the motor system in this frameworks’ scale-invariant outputs (i.e. modules-within-modules). Finally, our findings also highlight the crucial role of the individual muscle level in movement organization and promote further investigations on modular control that integrate scale.

To summarize the insights gained from applying the framework to benchmark datasets:

- The highly nested functional architecture of the muscle networks suggests human motor control is simplified via mechanisms mirroring dynamic programming (i.e. recursive decomposition of task demands across hierarchically structured modules).
- Coarse-grained control mechanisms correlate with poorer balance performance at the inter-muscular level but with better balancing intramuscularly.
- The gamma frequency band can explain much of the task-relevant information found among lower frequency-bands during balance control.
- Motor variability in older adults has a functional underpinning characterized by concurrent increases and decreases in functional integration at inter- and intra-muscular scales respectively.
- Independent muscle control is related to better balance performance but also with older age during a reach-and-grasp task.

In conclusion, we have developed and successfully applied a computational framework for the extraction of functionally diverse muscular interactions across multiple scales. Our approach provides a more detailed and precise account of the functional organization of the motor system by introducing a more nuanced and generalizable definition of muscles working together which, consequently, has direct benefits in the clinical setting and in engineering applications (e.g. predicting motor intention). We were able to comprehensively characterize the functional underpinnings of several distinct motor tasks while ensuring physiological relevance and generalizability. This principled aligns current approaches to muscle synergy analysis with the forefront of theoretical work on movement modularity, offering improved flexibility and opportunities to future investigations through nuanced perspectives on movement control.

## Limitations of study

The interactions quantified here do not imply a causal relationship to behavior, and so their direct effects on motor task performance remains a current limitation of the framework, the outputs of which should only be interpreted as different types of muscle-task statistical relationships. This is important as established research on muscle synergies has provided causal links between the summation of individual muscle activities within a synergy and force outputs ^61^, hence specific efforts towards aligning the presented framework with this work should be implemented. In future work, we will use multilevel neural interactions to predict motor intention and execution and perturb them using stimulation techniques to reveal their causal roles in motor behavior. Finally, as the exact neural substrates underpinning the different types of functional muscle interaction are currently not known, in future work we also aim to formally identify them by quantifying cortico-muscular interactions and motoneuron level modules using this framework.

## Resource availability

## Lead contact

Further information and requests for resources should be directed to and will be fulfilled by the lead contact, Dr. David O’ Reilly.

## Materials availability

This study did not generate reagents.

## Data and code availability

- Data will be shared by the lead contact upon reasonable request.
- Code and example data is available at the following repository: https://github.com/DelisLab/Muscle_PID.
- Any additional information required to re-analyze the data reported in this paper is available from the lead contact upon request.

## Acknowledgments

We would like to thank Daniel Chicharro for the helpful discussions on Partial Information Decompositions. This study was funded by the Biotechnology and Biological Sciences Research Council (BBSRC).

## Author contributions

DOR: Methodological development, conceptual development, coding, testing, data curation (Dataset 2), processing and framework application, manuscript writing and editing.

WS: Data curation (Dataset 3).

PH: Data curation (Dataset 1), manuscript review.

RA: Data curation (Dataset 2), Data pre-processing, manuscript editing. SA: Data curation (Dataset 3), manuscript review.

ID: Supervision, conceptual and methodological development, manuscript review and editing.

## STAR Methods

### Experimental Model and Study Participant Details

#### Data acquisition and experimental conditions

To illustrate our framework, we applied it to three datasets of EMG signals and corresponding continuous task parameters recorded while human participants performed different motor tasks. In dataset 1 (Fig.4(a)) ^38^, 3 healthy, adult participants performed whole-body, unimanual point-to-point reaching movements in various directions and to varying heights while EMG from 30 muscles (tibialis anterior, soleus, peroneus, gastrocnemius, vastus lateralis, rectus femoris, biceps femoris, gluteus maximus, erector spinae, pectoralis major, trapezius, anterior deltoid, posterior deltoid, biceps and triceps brachii) across both hemibodies were captured (Fig.(a)). Alongside these EMG recordings, 3D kinematic data from 18 body locations (elbow, wrist, mid-arm, index finger, shoulder, hip, knee, ankle and foot) across both hemibodies were captured along with additional kinematics from the head, right eye, left ear, and the center of pressure. Participants performed ∼2160 pseudo randomized trials each in structured blocks across two days to avoid fatigue. Movement onsets and offsets were determined at the timepoints which the index finger kinematic was 5% above and below its peak velocity in the trial respectively. Ethical approval was given as detailed in the parent paper ^38^.

In dataset 2 (Fig.4(b)), 3 healthy participants performed 10 consecutive trials of balancing on a balance board (Model 16130 Stability Platform, Lafayette Instrument) while self-induced perturbations were experienced along the frontal plane. Each trial lasted 30 seconds in which participants were instructed to maintain a balance board position parallel to the floor as best they could while focusing ahead at eye-level on a dot on the wall (<5 meters distance). Between trials, participants had 1 minute to rest. EMG recordings (Delsys Trigno™ wireless EMG, sampling frequency: 2000Hz) were taken from the bilateral medial gastrocnemius (MGN), tibialis anterior (TA), rectus femoris (RF) and biceps femoris (BF) while the horizontal angular displacement of the balance board was simultaneously recorded. Ethical approval was given by the Faculty of Biological Sciences Ethical Review Committee, University of Leeds.

For dataset 3 (Fig.4(c)) ^39^, 14 young adults (22.1±2.4 years old, two left-handed, two males) and 18 older adults (71.6±6.9 years old, two left-handed, 8 males) performed a bimanual grasp-lift-hold-replace task of both a light (0.2kg) and heavy (0.4kg) object (two manipulanda made from carbon-filled nylon) while in a seated position in front of a table. The objects were placed on the table 75% of shoulder width and 70% of maximum reach for each participant who were instructed to grasp the object and lift it to a target height in front of them (300mm height) and to hold the object as still as possible at this position for 10 seconds. Following this holding period, the participant was instructed to replace the object(s) back on the starting position markers. Participants performed 10 consecutive repetitions for each weight condition while EMG signals from the bilateral anterior deltoid (AD), extensor carpi radialis (ECR), flexor carpi radialis (FCR) and abductor pollicis brevis (APB) were recorded (Delsys Trigno™, sampling frequency: 2000Hz). Grip forces were recorded from 50N load cells (Omega, LCM201-50), acquired from a 16-bit data acquisition card (National Instruments, USB-6002) and processed in Labview (v.1.4). For load forces, six degree-of-freedom models were created in Qualisys for each object and were used to compute their 3D acceleration from which net load forces were calculated with respect to object mass. Trials commenced from 100ms prior to first contact until 100ms after last point of contact of either hand with the object, determined via recorded kinematic data of the objects position. This research was approved by the Research Ethics Committee of the Faculty of Biological Sciences of University of Leeds and all methods conformed to the Declaration of Helsinki and were carried out in accordance with the University’s regulations. Written informed consent was obtained by all participants following guidelines of the University of Leeds.

### Data pre-processing

#### Intermuscular analyses

The processing of EMG signals from dataset 1 and 2 for the purpose of analyses in the intermuscular space included the application of a bidirectional low-pass Butterworth filter with zero-phase distortion (order: 4^th^, cut-off: 20Hz) to the rectified signals. Extrapolation of the kinematic data for dataset 1 and 2 to align with the corresponding EMG signals was carried out using a cubic spline method. For dataset 3, the signals were processed as described in ^39^. More specifically, the rectified EMG signals were low-pass filtered (filter: 4^th^ order Butterworth with zero-phase distortion, cut-off: 10Hz) and down-sampled to 200Hz to align with kinetic and kinematic datapoints. The filtered EMG signals were then normalized against their peak amplitudes across trials. The grip and load forces were smoothed using a low-pass filter (cut-off: 12Hz, filter: 4^th^ order Butterworth).

#### Intramuscular analyses

The processing of EMG signals for the purpose of intramuscular analyses in the intramuscular space were uniform across datasets 1-3. This processing firstly included the filtration of the raw, unrectified EMG signals into specific frequency bands (Delta [0.1-4 Hz], Theta [4-8 Hz], Alpha [8-12 Hz], Beta [12-30 Hz], Low-Gamma (Piper rhythm) [30-60 Hz], High Gamma (Gamma) [60-80 Hz]) using a low- and high-pass filter combination (bi-directional 4^th^ order Butterworth filters with zero-phase distortion). Then the absolute values from a Hilbert transform of the filtered signals representing their oscillatory amplitudes were extracted for further analysis. In the case of dataset 3, no down-sampling of the EMG signals occurred as carried out for intermuscular analyses. The task parameters for datasets 1-3 were all extrapolated using a cubic spline method to temporally align with the corresponding, processed EMGs and no further processing was carried out.

### Methods details

#### Quantifying functionally diverse muscular interactions

To decompose the information a pair of muscles [*m*_*x*_, *m*_*y*_] (or frequencies-specific amplitudes [*f*_*x*_, *f*_*y*_]) carries about *τ* into redundant, synergistic and unique components, we implemented the PID framework ^29,62,63^. PID stems from a related information-theoretic measure known as co-information (co-I) that, by contrasting the sum of shared task information in *m*_*x*_ and *m*_*y*_ each alone (*I*(*m*_*x*_; *τ*) + *I*(*m*_*y*_; *τ*)) against their joint task information (*I*(*m*_*x*_, *m*_*y*_; *τ*)), quantifies their multivariate mutual information (*II*(*m*_*x*_; *m*_*y*_; *τ*)) (Equation 1.1) ^28^. co-I results in a single value representing either a net redundancy (negative co-I values) or net synergy (positive co-I values) across the system. PID builds on this by removing the conflation of redundancy and synergy evident in co-I through the decomposition of *I*(*m*_*x*_, *m*_*y*_; *τ*) (i.e. the JMI) into separate redundant (*R*(*m*_*x*_: *m*_*y*_; *τ*)) and synergistic (*S*(*m*_*x*_: *m*_*y*_; *τ*)) information atoms and the unique task information provided by *m*_*x*_ (*U*(*m*_*x*_: *τ*|*m*_*y*_)) and *m*_*y*_ (*U*(*m*_*y*_: *τ*|*m*_*x*_)) (Equation 1.2).

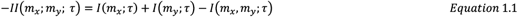

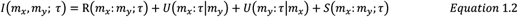

To perform PID here, we implemented a recent PID framework for Gaussian variables based on local information estimates that has proven useful in neuroscientific applications ^29,43,64–66^. We generated a multiplexed view of the muscular interactions underlying human movement by applying this method to all unique [*m*_*x*_, *m*_*y*_] (or [*f*_*x*_, *f*_*y*_]) and *τ* combinations for each participant. To make these separate computations directly comparable, we normalized each PID component by their collective sum total which is equal to the JMI ^43^. The resulting redundancy, synergy and two unique information estimates collectively form four symmetric adjacency matrices (*A*) (i.e. *A*^*T*^*A* = *I*) that represent the functionally similar, complementary or unique connectivities between muscles (frequencies) with respect to *τ*. When repeated across all available task variables τ and participants, *A* is a multiplex network of dimension [No.of [*m*_*x*_, *m*_*y*_] x [No.of τ x No.of participants]]. Thus, by applying network-theoretic statistical tools to *A*, we can identify functional modules carrying the same type of task information (i.e. redundant, synergistic, unique) (fig.1.2).

#### Identifying significant muscle interactions and their modular structure

The percolation threshold (*P*_*c*_) (i.e. a critical value specifying the probability of node connectivities occurring and at which large clusters and long-range connectivity begin to appear across a given network ^67^) has proven to be a fundamental constraint in nervous system organization ^1,6,68^. To identify the network connections in our framework that align with *P*_*c*_, we employed a modified percolation analysis ^6^. We applied this method to each layer of *A*, sparsifying the network with respect to the *P*_*c*_ expected from equivalently sized random networks by thresholding connectivities iteratively until the largest cluster in the network (the ‘*giant component’*) begins to be affected (Fig.3(C)).

Following this, in previous applications of the NIF ^19,30^, to determine the optimal number of clusters to extract with dimensionality reduction, we have implemented community detection protocols suitable for multiplexed networks ^34,69,70^. In the current study however, due to the highly nested structure found in *A*, just a single cluster was identified using these established methods. Therefore, to determine the optimal modular structure within a multiplexed network of muscles with overlapping functional affiliations, we employed a link-based community detection protocol ^35^. Specifically, we applied single-linkage hierarchical clustering to each layer of *A*, building dendrograms for each network layer describing the clustering of dependencies that is cut at a threshold determined by the maximal partition density (*D*) (Fig.3(D)), defined for a given partition of *M* links and *N* nodes into *C* subsets (Equation 2.1) ^35^. Here, *D* is the average of the number of links in a subset (*m*_*c*_) normalized by the possible maximum and minimum number of nodes with respect to the number of nodes those links touch (*n*_*c*_). This computation essentially unravels the nested network structure of each layer in *A*, resulting in a set of binary adjacency matrices that represent whether a muscle belongs to an identified cluster or not.

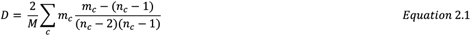

We then sum across all of these computed matrices across all layers of *A*, resulting in a single aggregate graph. Finally, we then apply the conventional community detection method based on a modularity maximization cost-function known as the Q-statistic (*Q*) (Equation 2.2) ^34,41^. For a given partition of the network, the Q-statistic compares the number of edges between node *i* and *j* (*A*_*ij*_) and what would be expected from an equivalent random network (*P*_*ij*_). In letting δ(*g*_*i*_, *g*_*j*_)=1 if nodes *i* and *j* belong to the same group (*g*) and 0 otherwise, thus this measure penalises partitions with a low ratio of within vs without cluster dependencies ^34,71^. The optimal cluster count to extract was defined as the partition that maximises this Q-statistic.

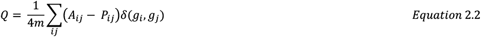

#### Extraction of low-dimensional motor components

Using the optimal cluster count derived a priori as the input parameter into dimensionality reduction, namely PNMF ^37^, we extracted low-dimensional components of motor behavior from muscle interactions of a specific type (i.e. redundant, synergistic or unique) across tasks and participants (Fig.3(E)). In the case of the intermuscular space, this input matrix (*A*) was of shape [No. of [*m*_*x*_, *m*_*y*_] x [No. of *τ* x No. of Participants]], while for the intramuscular space, *A* had a dimensionality of [No. of [*f*_*x*_, *f*_*y*_] x [No. of muscles x No. of *τ* x No. of Participants]]. As described in equation 3.1 for the *j*th module and single participant and task case, *A* is factorized into two components, ***v*** a vector of muscle weightings (*m*) of length equal to the number of unique muscle pairs (*K*) and corresponding activation coefficients (*s*). The extraction of the modules identified during model-rank specification is verifiable as shown in previous NIF applications ^30^. We note here that for the aim of extracting motor components associated with differences in the sampled population (see ‘*Hierarchies of functional muscle interactions capture distinct motor features’* in the results section), we input the normalized PID values into dimensionality reduction. However, for the aim of extracting generalizable motor components (see ‘*Generalizable components of functionally diverse muscular interactions’* and ‘*Functional modularity is scale-invariant across the human motor system’* of the results section), we input the non-normalized version of *A* into dimensionality reduction.

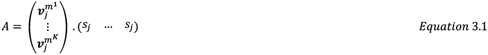

### Quantification and statistical analysis

#### Examining the generalizability of extracted motor components

To determine the generalizability of the extracted motor components, using Pearson’s correlation we determined the similarity between the extracted motor components and equivalent components extracted from a subset of the input data (i.e. when an individual participant or task was removed). We carried out this procedure for all tasks and participants and then focused our analysis on the correlation between functionally equivalent motor components (>0.5 correlation). To summarize this comparison, we converted the remaining coefficients into Fisher’s Z values, computed averages and standard deviations, and then reverted these values back to correlation coefficients.

#### Salient features of motor performance

To probe how functionally diverse inter- and intra-muscular interactions represent motor performance, we quantified ∑Error in specific ways for dataset 2 and 3 for use in separate statistical analyses.

For dataset 2, ∑Error was defined as the absolute cumulative error of the balance board parameter (|σ|) across the *n*th trial (Equation 4.1). σ was any deviation from 0 degrees of the balance board along the horizontal plane. Using a repeated measures correlation ^44^, associations between ∑Error and trial-specific activation coefficients from inter- and intra-muscular components were determined. For the intramuscular activation coefficients specifically, we averaged them across muscles to get a [No. of trials x No. of participants] size vector equivalent to the intermuscular coefficients.

For dataset 3, a binary variable representing participants’ age group (Young=0 vs. Old=1) was used as the dependent variable in a binary logistic regression model. ∑Error, a measure of motor variability, was used as the predictors in this model. ∑Error was defined as the absolute cumulative sum across trials of an inter-or intra-muscular activation coefficient σ demeaned with respect to their condition-specific average (i.e. light vs heavy objects) (Equation 4.1).

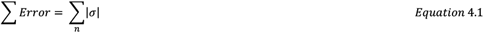

We calculated ∑Error of all inter- and intra-muscular components and input these as predictors of age group in the logistic regression model. An optimally parsimonious model, including the minimal number of predictors, was determined using forward selection via the Wald’s test criterion (inclusion: p<0.05, exclusion: p>0.1).

#### Subnetwork analysis

To illustrate the relative importance of individual muscles on the depicted human body models ^40^, we determined the total communicability (*C*(*i*)) of individual nodes (*i*) ^42^. *C*(*i*) is defined as the row-wise sum of all matrix exponentials in the adjacency matrix (*A*) that consider the number of walks between each pair of nodes *i* and *j* (Equation 5.1) ^36,42^.

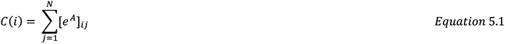

To emphasize salient functional connectivities present in the motor components, we further sparsified all dependencies with a below average network communicability and illustrated the output on the accompanying human body models ^36,40^. To uncover salient subnetwork structures consisting of more closely functionally related muscles (indicated by node color on the human body model), we applied the community detection algorithm described in equation 2.3 to the extracted motor components ^34,41^.

