## Supplementary materials for "Quantifying the diverse contributions of hierarchical muscle interactions to motor function"

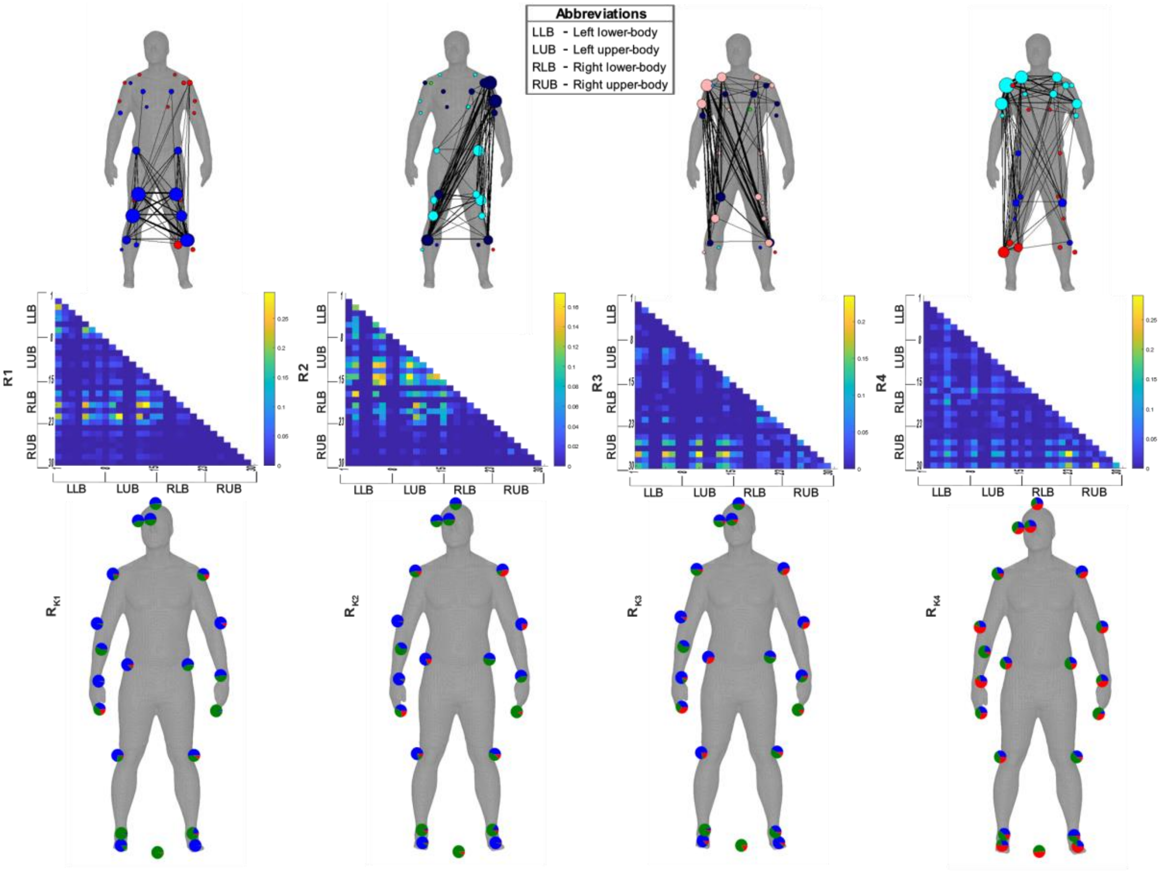


**Supp. Fig.1**: The $R$ muscle networks (**R1-R4**) extracted from the three participants of dataset 1 who performed a variety of whole-body reaching movements. Above R1-R4, their accompanying human body models with muscle networks overlaid illustrating the strongest connectivities (edge-width) [37], the subnetwork community structure (node color) and network centrality (relative node size) [32,38,39]. Below the adjacency matrices, the proportional encoding of XYZ kinematics in the activation coefficients averaged across participants ($\boldsymbol{R_{K1}}$-$\boldsymbol{R_{K4}}$) are depicted as pie charts (anteroposterior (red), mediolateral (green) and vertical (blue) directions) on corresponding bodily locations of a human body model.

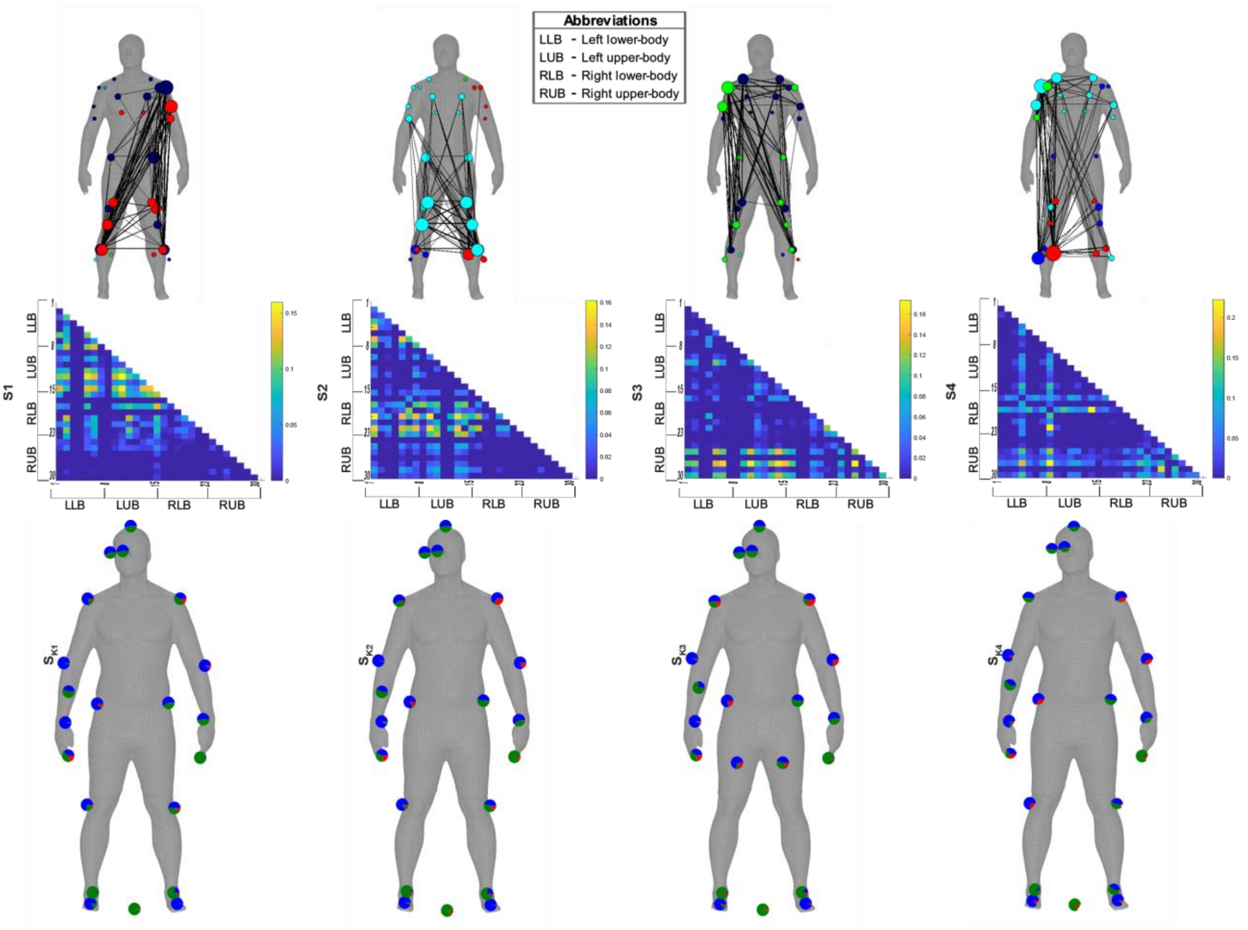


**Supp. Fig.2**: The $S$ muscle networks (**S1-S4**) extracted from the three participants of dataset 1 who performed a variety of whole-body reaching movements. Above S1-S4, their accompanying human body models with muscle networks overlaid illustrating the strongest connectivities (edge-width) [37], the subnetwork community structure (node color) and network centrality (relative node size) [32,38,39]. Below, the proportional encoding of XYZ kinematics in the activation coefficients averaged across participants ($\boldsymbol{S_{K1}}$-$\boldsymbol{S_{K4}}$) are depicted as pie charts (anteroposterior (red), mediolateral (green) and vertical (blue) directions) on corresponding bodily locations of a human body model.

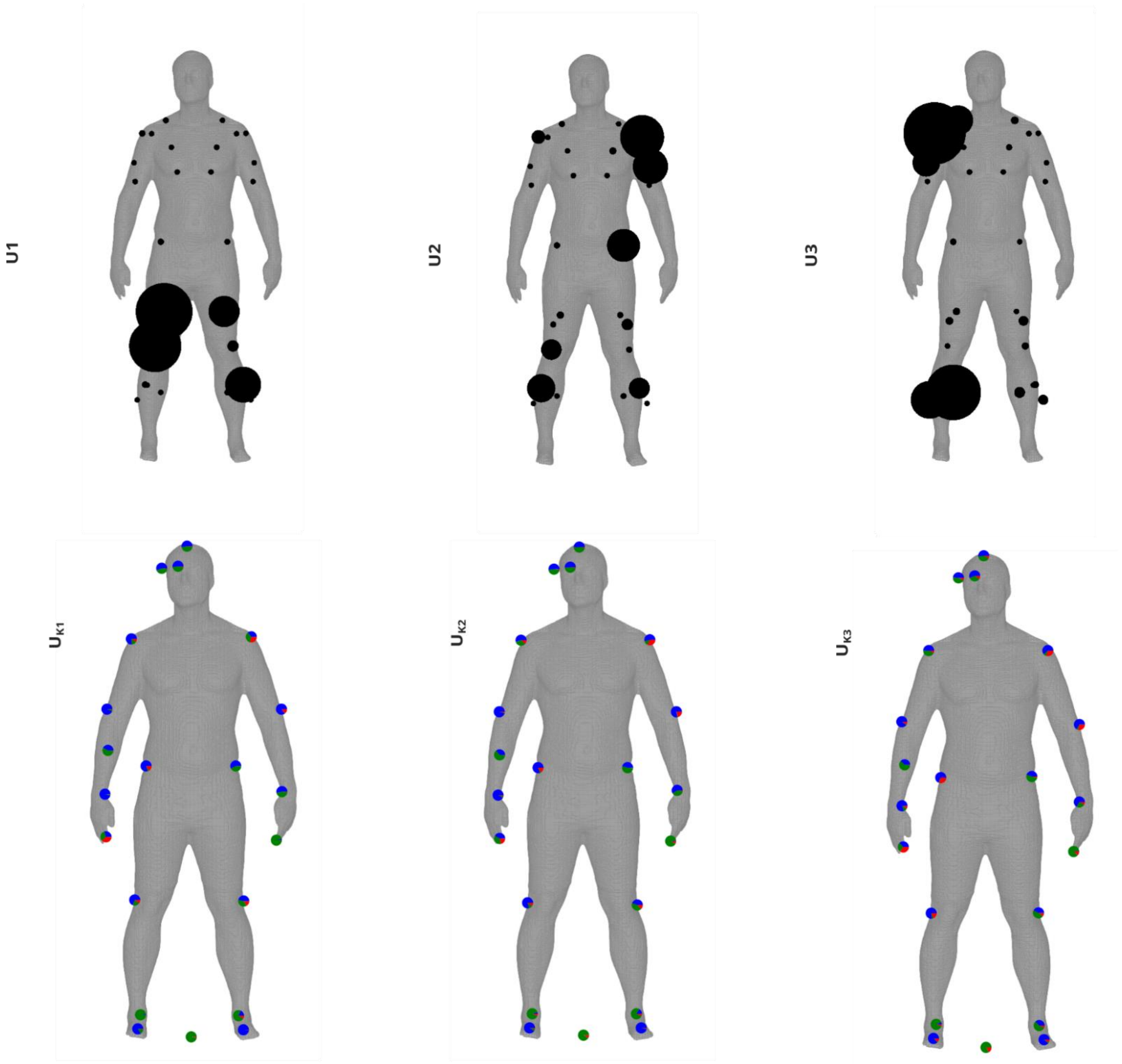


**Supp. Fig.3**: The $U_{xy}$ modules (**U1-U3**) extracted from the three participants of dataset 1 who performed a variety of whole-body reaching movements. The $U_x$ and $U_y$ terms are not considered as a muscle coupling, as they encode the task information present in one muscle that is not present in another and vice versa, and so instead the average unique information ($U_{xy}$) for each muscle is presented as the relative side of black nodes on a human body model. Below, the proportional encoding of XYZ kinematics in the activation coefficients averaged across participants ($\boldsymbol{U_{K1}}$-$\boldsymbol{U_{K3}}$) are depicted as pie charts (anteroposterior (red), mediolateral (green) and vertical (blue) directions) on corresponding bodily locations of a human body model.

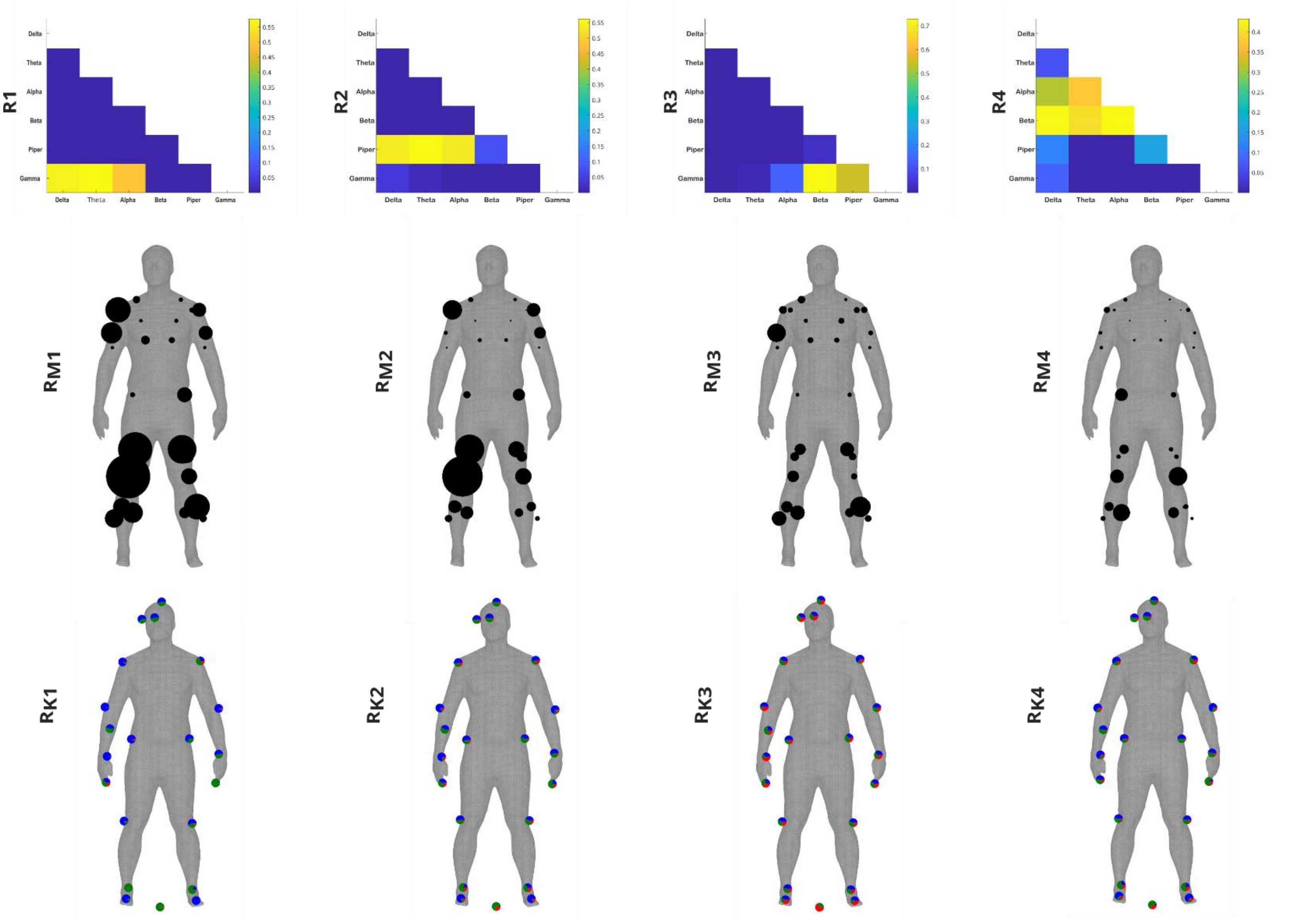


**Supp. Fig.4**: The $R$ intramuscular networks (**R1-R4**) extracted from the three participants of dataset 1 who performed a variety of whole-body reaching movements. Muscle- ($\boldsymbol{R_{M1}}$-$\boldsymbol{R_{M4}}$) and kinematic- ($\boldsymbol{R_{K1}}$-$\boldsymbol{R_{K4}}$) specific activation coefficients averaged across participants and each other are illustrated below their corresponding representations. For $\boldsymbol{R_{M1}}$-$\boldsymbol{R_{M4}}$, the relative size of the black nodes represents the activation of corresponding individual muscles. For $\boldsymbol{R_{K1}}$-$\boldsymbol{R_{K4}}$, intramuscular redundancy was determined with respect to 63 XYZ kinematic coordinates whose proportional task encodings are depicted as pie charts (anteroposterior (red), mediolateral (green) and vertical (blue) directions) on corresponding bodily locations of human body models.

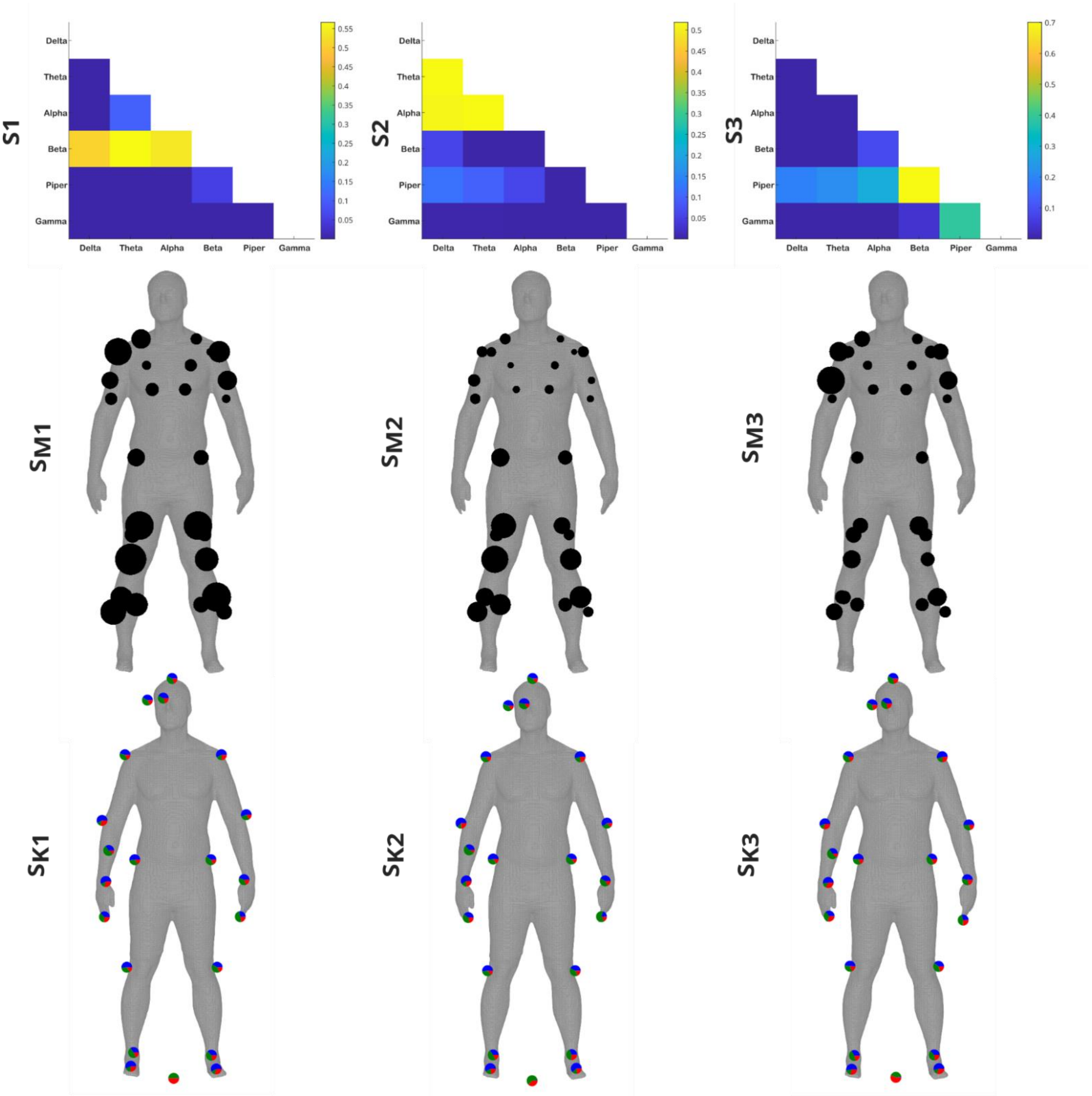


**Supp. Fig.5**: The $S$ intramuscular networks (**S1-S3**) extracted from the three participants of dataset 1 who performed a variety of whole-body reaching movements. Muscle- ($\boldsymbol{S_{M1}}$-$\boldsymbol{S_{M3}}$) and kinematic- ($\boldsymbol{S_{K1}}$-$\boldsymbol{S_{K3}}$) specific activation coefficients averaged across participants and each other are illustrated below their corresponding representations. For $\boldsymbol{S_{M1}}$-$\boldsymbol{S_{M3}}$, the relative size of the black nodes represents the activation of corresponding individual muscles. For $\boldsymbol{S_{K1}}$-$\boldsymbol{S_{K3}}$, intramuscular functional complementarity was determined with respect to 63 XYZ kinematic coordinates whose proportional task encodings are depicted as pie charts (anteroposterior (red), mediolateral (green) and vertical (blue) directions) on corresponding bodily locations of human body models.

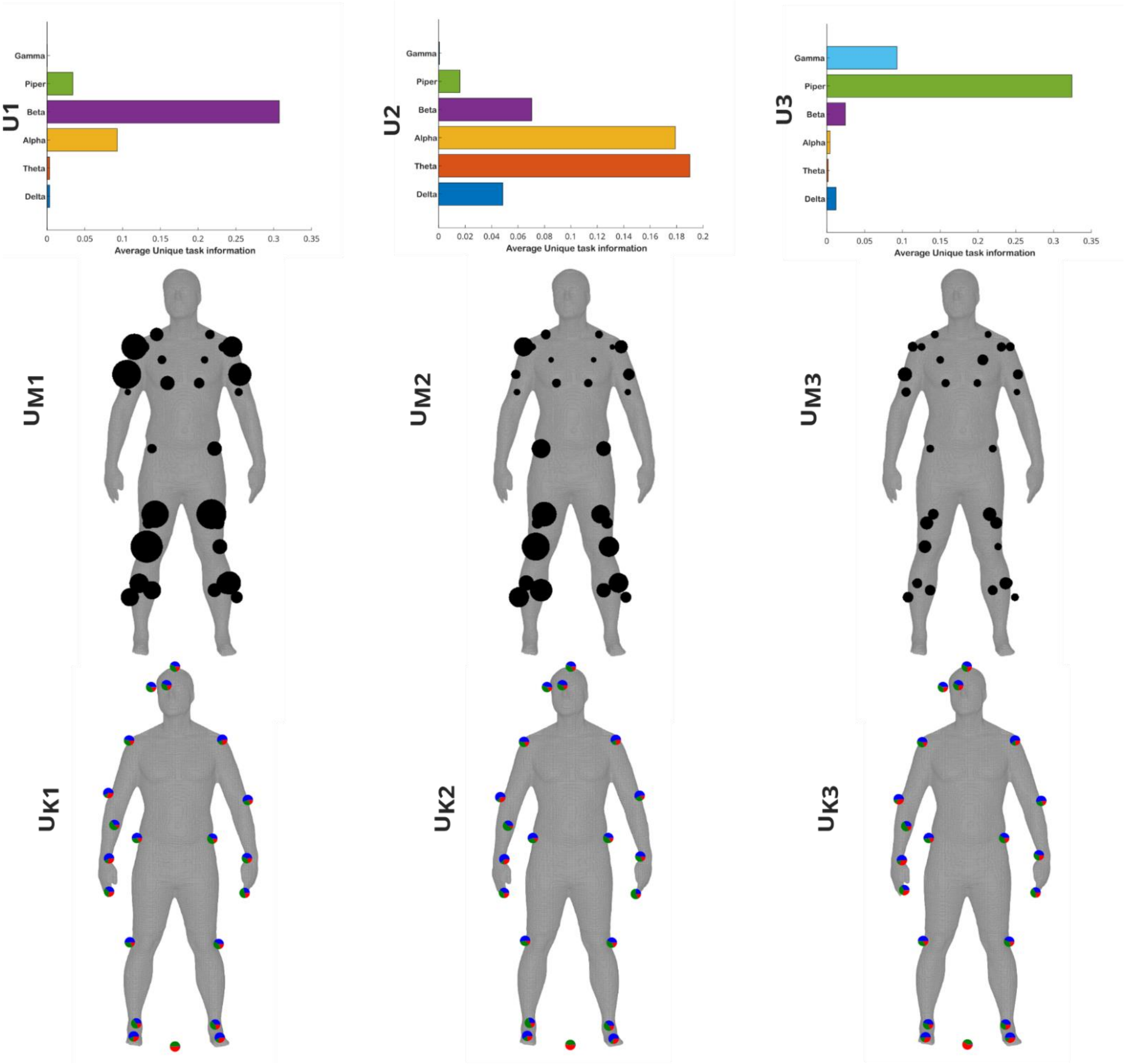


**Supp. Fig.6**: The $U_{xy}$ intramuscular networks (**U1-U3**) extracted from the three participants of dataset 1 who performed a variety of whole-body reaching movements. Muscle- ($\boldsymbol{U_{M1}}$-$\boldsymbol{U_{M3}}$) and kinematic- ($\boldsymbol{U_{K1}}$-$\boldsymbol{U_{K3}}$) specific activation coefficients averaged across participants and each other are illustrated below their corresponding representations. For $\boldsymbol{U_{M1}}$-$\boldsymbol{U_{M3}}$, the relative size of the black nodes represents the activation of corresponding individual muscles. For $\boldsymbol{U_{K1}}$-$\boldsymbol{U_{K3}}$, intramuscular functional independence was determined with respect to 63 XYZ kinematic coordinates whose proportional task encodings are depicted as pie charts (anteroposterior (red), mediolateral (green) and vertical (blue) directions) on corresponding bodily locations of human body models.
